# Patient-Calibrated Dynamical Modeling and Embedded Trend-Zone Predictive Control for Closed-Loop Deep Brain Stimulation in Parkinson’s Disease

**DOI:** 10.64898/2026.05.19.726196

**Authors:** Yu Fan, Lingxiao Guan, Yuxuan Wu, Xuesong Luo, Huiling Yu, Luming Li

**Affiliations:** National Engineering Research Center of Neuromodulation, School of Aerospace Engineering, Tsinghua University, Beijing, 100084, China; IDG/McGovern Institute for Brain Research, Tsinghua University, Beijing, 100084, China

**Keywords:** Deep brain stimulation, Predictive control, Parkinson’s disease, Closed loop systems

## Abstract

Closed-loop deep brain stimulation (cDBS) for Parkinson’s disease requires control strategies that tolerate noisy sensing, patient-specific stimulation responses, medication-related fluctuations, and embedded hardware constraints. We developed a patient-calibrated minute-scale dynamical model of subthalamic beta activity and an embedded explicit trend–zone predictive controller, eTZPC. The model combined a basal-ganglia mechanistic prior with stimulation-amplitude and medication-cycle recordings from five patients, and incorporated individualized stimulation-*β*_STN_ maps, fast- and slow-timescale stimulation responses, levodopa-related modulation, background drift, and observation noise.

eTZPC was designed to maintain *β*_STN_ activity within a patient-specific target zone under stimulation-amplitude, step-size, and quantization constraints. Compared with dual-threshold (DT) and proportional–integral–derivative (PID) controllers across four disturbance scenarios, eTZPC achieved target-zone regulation close to PID while reducing stimulation-switching burden toward the low-switching profile of DT. Ablation analyses identified distinct contributions of smoothing, trend prediction, patient-specific action modeling, and embedded explicit implementation. Parameter-mismatch tests showed that eTZPC was relatively robust to dynamic and disturbance-parameter deviations, but remained sensitive to errors in the steady-state stimulation–*β*_STN_ map. Patient-in-the-loop recordings in five patients further confirmed execution consistency and compliance with stimulation-boundary and step-size constraints. These findings support patient-calibrated dynamical modeling combined with low-complexity explicit control as a feasible framework for further embedded cDBS evaluation.

## I. Introduction

**D**eep brain stimulation (DBS) is an established neuromodulation therapy for advanced Parkinson’s disease (PD) [1], [2]. By delivering electrical pulses to the subthalamic nucleus (STN) or the internal segment of the globus pallidus (GPi), DBS can effectively alleviate core motor symptoms, including bradykinesia, rigidity, and resting tremor [3]. With the development of implantable sensing technologies, closed-loop DBS (cDBS) has emerged as an important direction for next-generation neuromodulation [4]–[7]. In such systems, stimulation parameters can be dynamically adjusted according to symptom-related neural biomarkers, among which beta-band power in STN local field potentials (*β*_STN_) is commonly used as an important representation of pathological neural activity in PD [8]–[13]. Recent clinical studies further suggest that long-term at-home cDBS based on embedded sensing–stimulation integrated devices is well tolerated and safe, and may reduce total electrical energy delivered (TEED) while maintaining clinical benefit [14]–[16].

Despite the clear promise of cDBS, its control design still faces a problem that has not been sufficiently clarified. Unlike conventional open-loop DBS, chronic closed-loop modulation is not merely a matter of “adjusting stimulation according to biomarker changes”; rather, it requires online control of a time-varying, disturbed, and partially observable system under multiple practical constraints imposed by the implantable device hardware and embedded software [7], [17]. First, biomarker measurements during fully implanted long-term recordings are susceptible to stimulation artifacts and multisource noise, including electrocardiographic activity, movement artifacts, and environmental interference [1], [18]. Second, stimulation-induced modulation of brain networks is nonlinear, hysteretic, and highly variable across individuals, and is further modulated by medication state and slow processes such as baseline drift and adaptive plasticity [19], [20]. Addressing these intertwined challenges requires more than a real-time tracking strategy: it calls for a control-relevant representation of the patient as a plant—one that captures the relevant dynamics, disturbances, and constraints in a form suitable for controller design, benchmarking, and validation [17], [21].

Predictive-control approaches, including conventional MPC and deep-learning MPC, have recently been explored for closed-loop DBS [21], [22]; however, how such control principles can be connected to patient-calibrated stimulation– *β*_STN_ dynamics, medication-related modulation, embedded constraints, and patient-in-the-loop execution remains insufficiently established.

Based on this rationale, we first established a patient-calibrated dynamical model that captures *β*_STN_ activity as shaped by both internal dynamics and external influences. The model characterizes the steady-state stimulation–*β*_STN_ relationship and the associated transition dynamics, while explicitly incorporating medication-related modulation, long-term baseline drift, and stochastic measurement noise as principal disturbance sources [19], [23].

On this basis, we designed an embedded explicit trend–zone predictive controller (eTZPC) to satisfy safety constraints and embedded implementation requirements, and compared it with conventional dual-threshold (DT) control and proportional–integral–derivative (PID) control [11], [24]. In addition, by comparing the full trend–zone predictive control (TZPC) with eTZPC, we evaluated whether embedded reduction preserved the main closed-loop performance [25], [26]. This study addressed two questions: first, what are the essential components of a control-relevant model for closed-loop DBS in PD, given the constraints visible in real patient data; and second, how do candidate closed-loop control strategies, including DT, PID, and predictive control, compare in robustness, constraint satisfaction, and engineering deployability under this model. Our results show that this modeling framework provides a unified basis for the control design and evaluation of closed-loop DBS, and supports feasibility validation of deployable control strategies under multiple realistic disturbance conditions.

## II. Methods

### A. Patient-Calibrated β_STN_ Dynamical Model

The *β*_STN_ dynamical model was organized into three components: a patient-specific fixed-stimulation steady-state map, 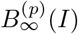 a fast–slow transition process after stimulation-amplitude changes; and medication/background modulation with observation-level noise for non-stimulation-related variability.

#### 1) Steady-State Stimulation–β_STN_ Relationship Calibrated by a Computational Neural Prior

We first established the steady-state stimulation–*β*_STN_ map *B*_*∞*_(*I*), which describes the *β*_STN_ level approached under a fixed stimulation amplitude *I*. Because the stimulation–*β*_STN_ relationship may be nonlinear and saturating, and its parameters vary substantially across patients, we constructed patient-specific maps by combining a mechanistic prior with data-driven calibration, rather than relying solely on generic parametric fitting of limited clinical data.

We established a basal-ganglia circuit model based on the Rubin–Terman framework [21], [27], [28] to generate a mechanistically constrained prior curve for *B*_*∞*_(*I*). The model included the STN, GPe, GPi, and thalamus, and DBS input to STN neurons was modeled through an electric-field propagation model. The complete neuronal equations, network connectivity, stimulation-field model, and parameter settings are provided in Supplementary Methods S1. This computational neural model was used as a shape prior for the stimulation–*β*_STN_ steady-state relationship, not as the online plant for closed-loop control.

To approximate clinical bipolar STN LFP recordings, the simulated LFP was computed as a differential volume-conduction signal from synaptic currents at the two recording contacts:

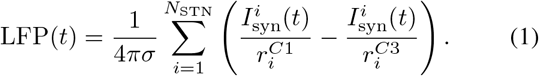

Here, *σ* denotes tissue conductivity and was set to 0.27 mS*/*mm [21], 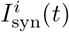 denotes the synaptic current of the *i*-th STN neuron, and 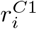 and 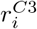 denote the distances from this neuron to the two bipolar recording contacts, respectively. Simulations were then performed at different stimulation amplitudes, and beta-band power was calculated from the bipolar LFP. The mean *β*_STN_ curve across simulations was defined as:

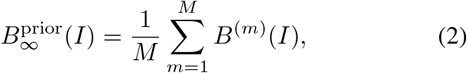

where *M* = 100, and *B*^(*m*)^(*I*) denotes the *β*_STN_ obtained in the *m*-th simulation under stimulation amplitude *I*. This curve was used to characterize the stimulation–*β*_STN_ steady-state shape generated by the mechanistic model.

After obtaining the mechanistic prior curve, this study further calibrated it using real patient stimulation–response data. Specifically, 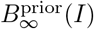 was fitted to the measured *β*_STN_ data of each patient through parameterized transformations, yielding the patient-specific steady-state map:

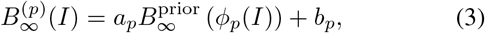

where *p* indexes patients, *ϕ*_*p*_(*I*) aligns the clinical stimulation-amplitude axis to the prior-curve coordinate, and *a*_*p*_ and *b*_*p*_ adjust the amplitude and baseline of the curve. Thus, 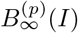 retains the nonlinear shape constraint of the mechanistic prior while incorporating patient-specific stimulation–*β*_STN_ scaling.

#### 2) Fast–Slow β_STN_ Dynamics with Adjustment of Stimulation Amplitudes

The steady-state map 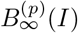 defines the *β*_STN_ level approached under fixed stimulation, but does not describe the transient response with stimulation changes. We therefore introduced a fast–slow dynamical formulation to capture the immediate and minute-scale components [21], [29] of the transition toward the new steady state.

Specifically, the post-switching *β*_STN_ response was decomposed into an immediate steady-state-dependent component and a slow relaxation component. The slow state *B*_*s*_(*t*) evolves toward the patient-specific steady-state level 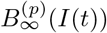:

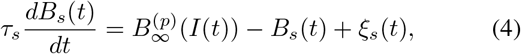

where *τ*_*s*_ is the relaxation time constant and *ξ*_*s*_(*t*) represents unmodeled endogenous fluctuation. The stimulation-related *β*_STN_ component was then expressed as a weighted sum of the current steady-state term and *B*_*s*_(*t*):

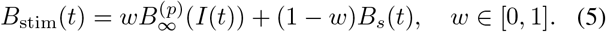

Here, *w* is the fast-component weight. After stimulation switching, 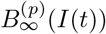changes with the new amplitude, whereas *B*_*s*_(*t*) relaxes toward the new steady state. This structure captures both the immediate response and the wash-in/wash-out dynamics without assuming full instantaneous steady-state replacement.

#### 3) Levodopa-Related Modulation and Background Drift

In addition to stimulation-driven fast–slow dynamics, chronic *β*_STN_ recordings contain non-stimulation-related variability arising from anti-parkinsonian medication, represented here by levodopa, and spontaneous brain-state fluctuations across arousal, behavioral, and circadian dimensions [19], [23], [29], [30]. These effects were simplified as changes:

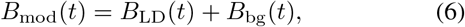

where *B*_LD_(*t*) denotes levodopa-related modulation and *B*_bg_(*t*) denotes slow background drift.

Combined with the stimulation-related component *B*_stim_(*t*) defined in Section 1.2, the latent *β*_STN_ state generated by the model without observation noise was defined as:

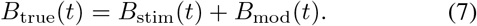

Here, *B*_true_(*t*) denotes the latent *β*_STN_ state generated by the model under the specified stimulation and background conditions. The controller-visible biomarker further included observation noise:

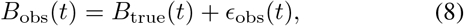

where *ϵ*_obs_(*t*) denotes observation-level disturbances, including LFP measurement noise, stimulation artifacts, electrocardiographic and movement artifacts, and spectral-estimation errors.

Levodopa-related modulation was estimated from medication-cycle recordings. Using medication intake as time zero, systematic *β*_STN_ changes were summarized as a normalized temporal profile *L*(*t*), and the patient-specific modulation term was defined as:

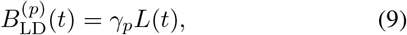

where *γ*_*p*_ denotes the patient-specific modulation amplitude. Estimation details and parameter values are provided in Supplementary Methods S4 and Tables S9–S10.

The amplitude and rate of *B*_bg_(*t*) were constrained by baseline drift observed in chronic recordings.

The model therefore separated non-stimulation-related variability into latent-state modulation and observation-channel noise. Closed-loop controllers updated stimulation using only *B*_obs_(*t*), whereas performance metrics were evaluated on the latent state *B*_true_(*t*).

#### 4) Real Data Sources and Parameter Calibration

Recordings from five patients with PD were used to calibrate the dynamical model. Calibration followed the model hierarchy: fixed-amplitude stimulation recordings were used to fit the patient-specific steady-state map 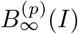 stimulation-switching recordings were used to estimate the fast-component weight *w*_*p*_ and relaxation time constant *τ*_*s,p*_; and medication-cycle/chronic recordings were used to estimate levodopa modulation, background drift and observation-noise levels. The complete fitting procedures are provided in Supplementary Methods S3-S4, and individualized parameters are summarized in Table I and Supplementary Table S9.

**TABLE I:** Patient-specific model and simulation parameters.

| Parameter | P1 | P2 | P3 | P4 | P5 |
| --- | --- | --- | --- | --- | --- |
| $I_{\min}$ (mA) | 2.05 | 1.80 | 2.95 | 1.25 | 1.25 |
| $I_{\max}$ (mA) | 2.55 | 2.10 | 3.25 | 1.40 | 1.45 |
| $B_L$ | 0.8977 | 1.9447 | 1.4725 | 1.1761 | 1.0930 |
| $B_H$ | 0.9545 | 1.9903 | 1.5825 | 1.8022 | 1.9526 |
| $\gamma_p$ | 0.2546 | 0.2495 | 0.3051 | 1.6238 | 1.9455 |
| $\tau_s$ (min) | 18.2434 | 14.2526 | 22.9745 | 18.5714 | 19.6494 |
| $w_p$ | 0.0046 | 0.0321 | 0.0062 | 0.0688 | 0.1839 |
| $\sigma_\epsilon$ | 0.0462 | 0.0975 | 0.0858 | 0.0072 | 0.0051 |
| $\sigma_\xi$ | 0.0171 | 0.0545 | 0.0219 | 0.0367 | 0.0389 |

Note: $I_{\min}$ and $I_{\max}$ denote the lower and upper stimulation bounds used during closed-loop control. $[B_L, B_H]$ denotes the patient-specific target beta zone. $\gamma_p$ denotes the levodopa modulation amplitude. $\tau_s$ and $w_p$ denote the slow relaxation time constant and fast response weight. $\sigma_\epsilon$ and $\sigma_\xi$ denote observation-noise and endogenous fluctuation strengths, respectively.

### B. Closed-Loop Control Strategy

Based on the patient-calibrated model and the general predictive-control principle [25], we developed a single-step trend–zone predictive control (TZPC) strategy. The controller aimed to keep *β*_STN_ within a patient-specific target zone [*B*_*L*_, *B*_*H*_] under stimulation-amplitude, step-size, and quantization constraints, rather than tracking a single reference value. At each control step, TZPC used the smoothed beta level, local trend, and patient-specific stimulation–beta map 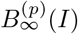 to select a constrained stimulation update by balancing zone violation, predicted beta change, and update cost.

For embedded deployment, the full TZPC action prediction was locally linearized within each patient’s clinical stimulation working range, yielding an explicit trend–zone feedback law, termed eTZPC. This reduced online computation to smoothing, trend estimation, zone-error calculation, gain-based updating, quantization, and boundary checking. Details are provided below.

#### 1) Trend–Zone Predictive Control Formulation

Let *B*_obs,*k*_ denote the noisy *β*_STN_ observation obtained from the implantable system at the *k*-th control step. To reduce the influence of observation noise, stimulation artifacts, and spectral-estimation errors on control decisions, the observed sequence was first recursively smoothed:

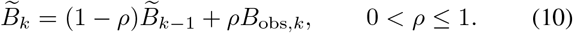

Here, 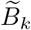 denotes the smoothed biomarker level, and *ρ* is the smoothing coefficient. The local trend was then estimated from the change between adjacent control cycles:

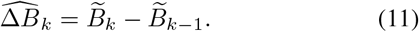

Without considering the effect of any newly applied stimulation action in the current control cycle, the no-action prediction of *β*_STN_ at the next step was defined as:

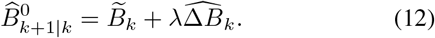

Here, *λ* is the look-ahead weight that controls how strongly the estimated trend is extrapolated into the next-step prediction.

Let *u*_*k*_ = Δ*I*_*k*_ denote the stimulation-amplitude update at the *k*-th control cycle. For the *p*-th patient, the predicted effect of a candidate stimulation action *u*_*k*_ on *β*_STN_ was given by the patient-specific steady-state map 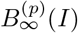:

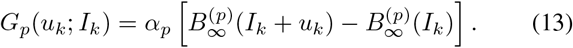

Here, *α*_*p*_ ∈ (0, 1] denotes the proportion of the steady-state difference that can be realized within one control cycle, thereby linking this formulation to the fast–slow dynamics described in Section 1. According to the fast–slow response model in Section 1.2, *α*_*p*_ can be approximately determined by the fast-component weight and the slow-process time constant:

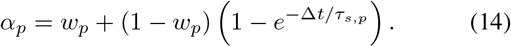

Thus, *G*_*p*_(*u*_*k*_; *I*_*k*_) represents the action effect after accounting for the response proportion achievable within the one minute’s update cycle.

The one-step predicted value after applying the current stimulation action was written as:

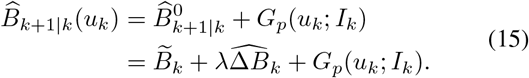

TZPC selects the stimulation update by minimizing the zone-predictive cost:

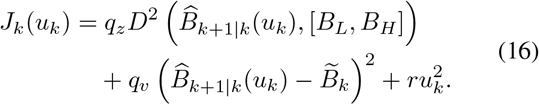

The first term penalizes violation of the target zone by the predicted *β*_STN_, the second term penalizes the change of the predicted value relative to the current level, and the third term penalizes the stimulation-amplitude update. The coefficients *q*_*z*_, *q*_*v*_, and *r* denote the corresponding penalty weights. The zone-violation function was defined as:

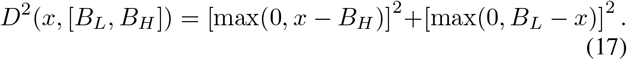

When the predicted value lies within the target zone, the zone-violation term is inactive, and the controller mainly acts as a trend-damping controller. When the predicted value exceeds the upper boundary or falls below the lower boundary, the zone-violation term is activated, and the controller automatically introduces a corrective action proportional to the zone-violation error.

The full theoretical action was given by:

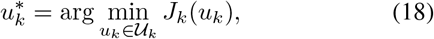

where the admissible action set was defined by the stimulation step-size and safety-bound constraints:

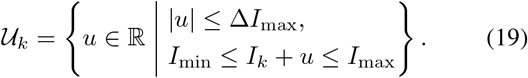

#### 2) Embedded Explicit Implementation of TZPC

During embedded online operation, stepwise computation of the nonlinear *G*_*p*_(*u*_*k*_; *I*_*k*_) and solution of the cost-minimization problem would increase implementation complexity. Therefore, within the clinically selected stimulation working range of each patient, the action effect was locally linearized offline as:

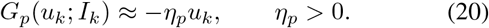

Here, *η*_*p*_ denotes the effective stimulation sensitivity of the *p*-th patient within the working stimulation range. This parameter was obtained by least-squares linear fitting of the patient-specific steady-state map 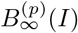 within the working range. If the fitted slope is ŝ _*p*_, then:

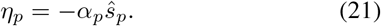

Under local linearization and fixed-gain constraints, TZPC was transformed into an explicit trend–zone feedback law by minimizing the optimization objective *J*_*k*_(*u*_*k*_) in (16). First, the no-action prediction error relative to the target zone was defined as:

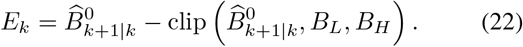

Here, clip(*x, a, b*) = min(max(*x, a*), *b*) denotes clipping *x* to the interval [*a, b*]. When 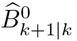 lies within the target zone, *E*_*k*_=0.it is below the lower boundary, *E*_*k*_ *>* 0.When it is above the upper boundary,*E*_*k*_ *<* 0.

The online stimulation-update command was written as:

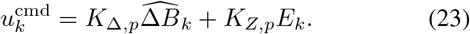

The first term is the trend-damping term, and the second term is the zone-correction term. The corresponding gains are:

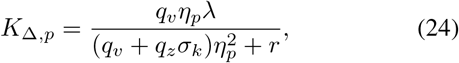

and

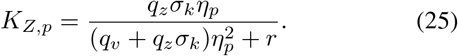

Here, *σ*_*k*_ indicates whether the zone-violation term is activated:

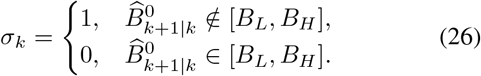

The detailed derivation is provided in the Supplementary Methods.

Finally, the control output was processed through the maximum step-size constraint, stimulator amplitude-resolution constraint, and stimulation safety boundaries:

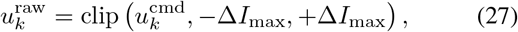

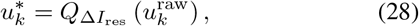

and

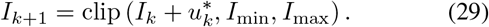

Here, 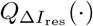 denotes quantization of the stimulation update to an integer multiple of the minimum amplitude resolution, with Δ*I*_res_ = 0.025 mA. Thus, online eTZPC can be regarded as an explicit implementation of full TZPC under local linearization, fixed gains, and embedded constraints. In this formulation, 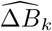 is responsible for anticipatory trend response, whereas *E*_*k*_ is responsible for target-zone correction.

A slightly inward-contracted control zone was used during online decision making to reduce quantization-induced dead zones; formal performance evaluation used the original target zone [*B*_*L*_, *B*_*H*_]. Details are provided in Supplementary Methods S5.

#### 3) Other Control Strategies

Two representative controllers were adopted for comparison: conventional dual-threshold (DT) control [11] and proportional–integral–derivative (PID) control [24], [31]. DT control represents a reactive zone-based strategy, whereas PID represents classical continuous error feedback. All controllers used the same target zone, stimulation bounds, maximum step size, amplitude resolution, and update interval.

The DT controller updated stimulation according to the smoothed beta level:

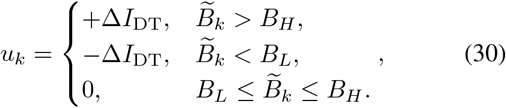

where Δ*I*_DT_ is the fixed update step.

The PID controller used the target-zone center as the reference:

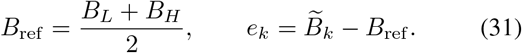

The commanded stimulation update was:

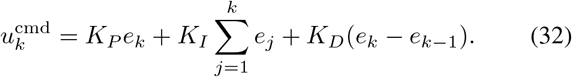

where *K*_*P*_, *K*_*I*_, and *K*_*D*_ are the proportional, integral, and derivative gains. The PID output was subjected to the same step-size, quantization, and stimulation-boundary constraints as eTZPC.

#### 4) Controller-Parameter Optimization

Controller parameters were selected using a predefined offline tuning procedure. To reduce performance overestimation, an additional LD + endogenous fluctuation simulation environment constructed from P1 was used only for parameter optimization and was excluded from all formal evaluations, disturbance-scenario comparisons, ablation analyses, and parameter-mismatch tests. The optimized parameters were *q*_*z*_, *q*_*v*_, *r*, and *λ* for eTZPC, and *K*_*P*_, *K*_*I*_, and *K*_*D*_ for PID. The DT update step was selected from {0.025, 0.05, 0.075, 0.10}mA. Continuous parameters were optimized using multi-start derivative-free pattern search. All selected parameters were fixed across patients, test scenarios, and disturbance conditions. The smoothing coefficient was fixed at *ρ* = 0.5 for all controllers with smoothing. Detailed tuning objectives and final parameter values are provided in the Supplementary Methods.

### C. Simulation and Evaluation Protocol

#### 1) Closed-Loop Simulation Settings

Closed-loop simulations were performed using the patient-calibrated *β*_STN_ model. For each patient, the plant was defined by the calibrated steady-state map 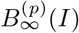, fast–slow dynamic parameters, levodopa modulation, background drift and observation noise. At each control step, the model generated the latent beta state 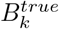 and the controller-visible biomarker:

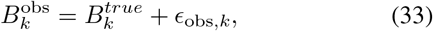

where *ϵ*_obs,*k*_ denotes observation-level disturbance. Controllers updated stimulation using only 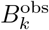 and its history, and all outputs were constrained by the same step-size limit, amplitude resolution, and safety boundaries.

The simulation update interval was set to Δ*t* = 1 min. Each simulation lasted 480 min, corresponding to an 8-h recording window that could cover the main rising, maintenance, and decay phases of the levodopa effect. Each patient–scenario– controller combination was repeated 10 times to evaluate the sensitivity of the results to random noise realizations and initial disturbance phases.

#### 2) Test Scenarios

Four test scenarios were defined to distinguish the effects of latent-state endogenous fluctuation and observation-channel noise on closed-loop control. Endogenous process disturbance acted on the latent *β*_STN_ dynamics, whereas observation noise acted only on the controller-visible biomarker 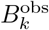.

In the **LD-only scenario**, the model included only the stimulation response and levodopa-related modulation, without additional endogenous process disturbance or observation noise.

In the **LD + endogenous fluctuation scenario**, Gaussian process disturbance was added to the latent slow-state dynamics:

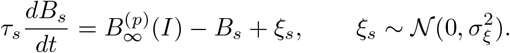

In the **LD + endogenous fluctuation + Gaussian observation noise scenario**, zero-mean Gaussian noise was further added to the observation channel:

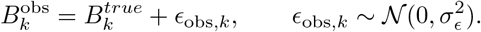

In the **LD + endogenous fluctuation + sinusoidal observation disturbance scenario**, a periodic observation disturbance was added:

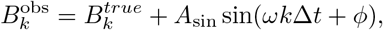

where *A*_sin_, *ω*, and *ϕ* denote the disturbance amplitude, angular frequency, and initial phase, respectively.

All scenarios used the same patient model, target zone, stimulation bounds, maximum step size, and update interval. Disturbance strengths were estimated from chronic-recording residuals after removing stimulation-and levodopa-related trends. Low-frequency innovations defined *σ*_*ξ*_, high-frequency components defined *σ*_*ϵ*_, and *A*_sin_ = 2.5*σ*_*ϵ*_ with a 480-min period. *B*_bg_(*t*) was not tested separately because it mainly represented slower circadian-scale drift.

Baseline comparisons were performed under all four scenarios using the metrics defined below.

#### 3) Progressive Ablation Analysis

Progressive ablation tested the incremental contribution of each eTZPC module. **A0** was current-value dual-threshold control; **A1** added observation smoothing; **A2** used the no-action prediction 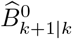 for threshold decisions; **A3** added patient-specific nonlinear action prediction and corresponded to full TZPC; and **A4** used the locally linearized explicit law, corresponding to eTZPC. All variants used the same plant, target zone, constraints, and metrics.

#### 4) Parameter-Mismatch Robustness Analysis

To evaluate robustness to model-calibration errors, parameter-mismatch tests were performed by perturbing the closed-loop plant while keeping all controller parameters fixed. Thus, the controller continued to use the originally calibrated patient model, whereas the simulated plant was altered to represent potential mismatch between offline calibration and true patient dynamics.

Four classes of mismatch were tested: steady-state stimulation–*β*_STN_ mapping mismatch, fast–slow stimulation-response mismatch, medication/background modulation mismatch, and noise-strength mismatch. These perturbations respectively represented errors in the estimated stimulation-response curve, stimulation-transition dynamics, slow non-stimulation-related modulation, and disturbance intensity. Closed-loop simulations were rerun under each perturbation condition, and the same performance metrics as in the baseline comparison were calculated, including target-zone time, zone-violation area, mean stimulation amplitude, and switching count.

#### 5) Control Performance Metrics

Closed-loop control performance was evaluated from four aspects: target-zone maintenance, zone-violation error, stimulation burden, and control smoothness. Because the control objective in this study was to maintain *β*_STN_ within the target zone rather than to track a single fixed reference value, zone-control metrics were used as the primary performance measures.

First, the time-in-range ratio was defined as:

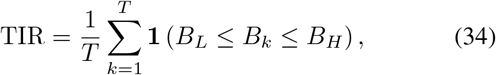

where *T* denotes the total number of simulation steps, and **1**(·) denotes the indicator function.

Second, the integrated zone absolute error was defined as:

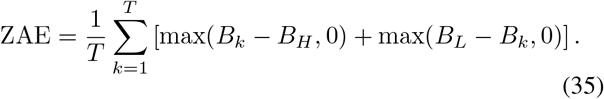

Third, to evaluate stimulation burden, the mean stimulation amplitude was calculated as:

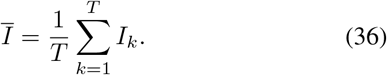

Fourth, to evaluate control smoothness, the number of stimulation adjustments was calculated as:

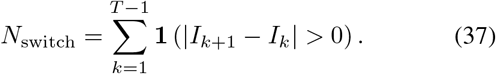

#### 6) Online Execution-Consistency Evaluation

For embedded validation, eTZPC was implemented in the implantable pulse generator and tested during *>*8-h online closed-loop recordings in all patients. Patients withheld dopaminergic medication for at least 12 h before recording and then followed their routine daytime medication schedule. Device-stored smoothed beta energy and stimulation amplitudes were exported to verify stimulation-boundary and step-size compliance. The recorded biomarker sequence was replayed offline using the same eTZPC law, and execution consistency was defined as the proportion of matched online and replayed updates. This analysis assessed embedded execution consistency rather than clinical efficacy.

#### 7) Statistical Analysis

Simulation metrics were averaged across repeated noise realizations to obtain one value per patient, scenario, and controller, and group results were reported as mean ±SD. Controller comparisons used paired patient-level Friedman tests followed by Holm–Bonferroni-corrected Wilcoxon signed-rank tests when appropriate. Ablation analyses used predefined adjacent paired comparisons; mismatch and online execution-consistency analyses were descriptive. Significance was set at *p <* 0.05.

## III. Results

### A. The Patient-Calibrated Dynamical Model Captured Stimulation- and Medication-Related Modulation of β_STN_

Basal ganglia network simulations first generated a mechanistic prior curve describing the relationship between stimulation amplitude and the steady-state level of *β*_STN_, showing that the simulated steady-state *β*_STN_ level decreased as DBS amplitude increased (Fig. 1A). This prior curve was then individually aligned to the stimulation-amplitude modulation recordings from five patients with PD, yielding patient-specific stimulation–*β*_STN_ steady-state maps (Fig. 1B). Fig. 1B shows a representative calibration result for P1. Across patients, the stimulation working range, *β*_STN_ target zone, stimulation sensitivity, and fitting error all showed marked interindividual differences, supporting the use of patient-calibrated plants in the subsequent closed-loop simulations. A summary of the patient-level calibration parameters is provided in Table I, and the complete fitted parameters are listed in Table S9.

**Fig. 1.**
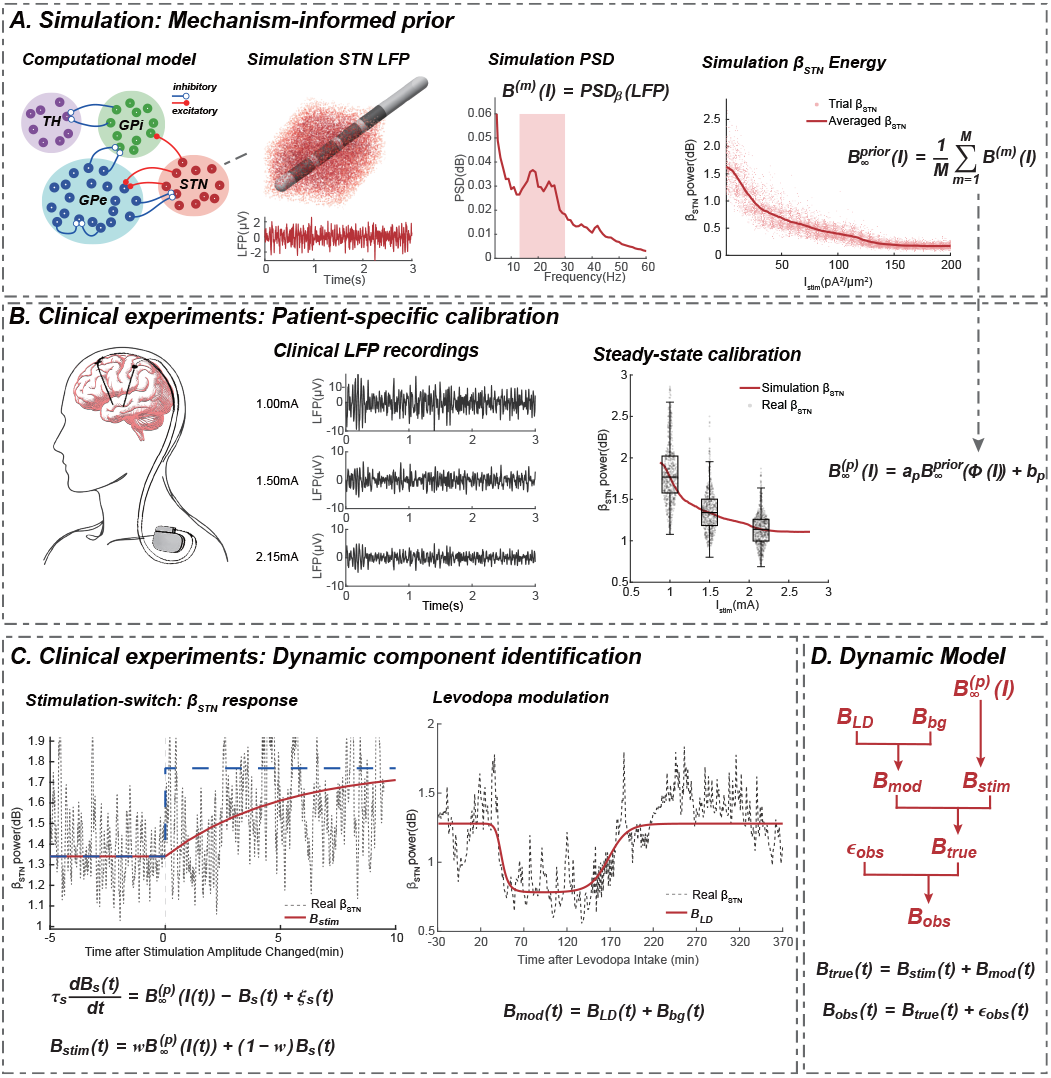
Patient-calibrated minute-scale ***β***_**STN**_ dynamical modeling framework. (A) Mechanism-informed prior generation from a basal-ganglia computational model. (B) Patient-specific calibration of the steady-state stimulation–***β***_**STN**_ map. (C) Identification of fast–slow stimulation dynamics and levodopa-related modulation from clinical recordings. (D) Integrated minute-scale model including stimulation-driven dynamics, background modulation, endogenous variability, and observation noise. DBS, deep brain stimulation; STN, subthalamic nucleus; LD, levodopa.

Beyond the steady-state mapping, stimulation-switching recordings further showed that *β*_STN_ did not reach the new steady state instantaneously after amplitude changes, but instead exhibited a fast response followed by a minute-scale relaxation process (Fig. 1C). Accordingly, the patient-specific model was extended with a fast–slow stimulation-response component to describe the transition dynamics after stimulation adjustment. Medication-cycle recordings further showed that *β*_STN_ was also influenced by levodopa-related slow-timescale modulation. This modulation was represented using a shared temporal profile across patients together with patient-specific modulation amplitudes (Fig. 1C). The shared levodopa temporal parameters are listed in Table S10, and the patient-specific modulation amplitudes are included in Table I and Table S9.

Finally, the patient-specific stimulation–*β*_STN_ steady-state mapping, fast–slow dynamics with stimulation switching, levodopa modulation, background drift, and observation noise were integrated into a unified *β*_STN_ dynamical model (Fig. 1D). This model defined the patient-calibrated plant used in the subsequent closed-loop controller comparison, disturbance-scenario testing, progressive ablation analysis, and parameter-mismatch robustness evaluation.

### B. eTZPC Transformed the Patient-Calibrated Model Into a Constrained Embedded Closed-Loop Control Law

Based on the patient-calibrated *β*_STN_ plant described above, this study further transformed the stimulation–*β*_STN_ dynamical model into a trend–zone predictive control framework for minute-scale cDBS (Fig. 2). Unlike conventional DT control, which triggers stimulation adjustment only after the currently observed *β*_STN_ exceeds the target boundary, this framework first smooths the observed *β*_STN_ and combines it with the local trend to generate a one-step no-action prediction (Fig. 2A). Thus, the control decision no longer depends only on whether the current *β*_STN_ is outside the target zone, but instead depends on whether the predicted *β*_STN_ is at risk of leaving the patient-specific target zone.

**Fig. 2.**
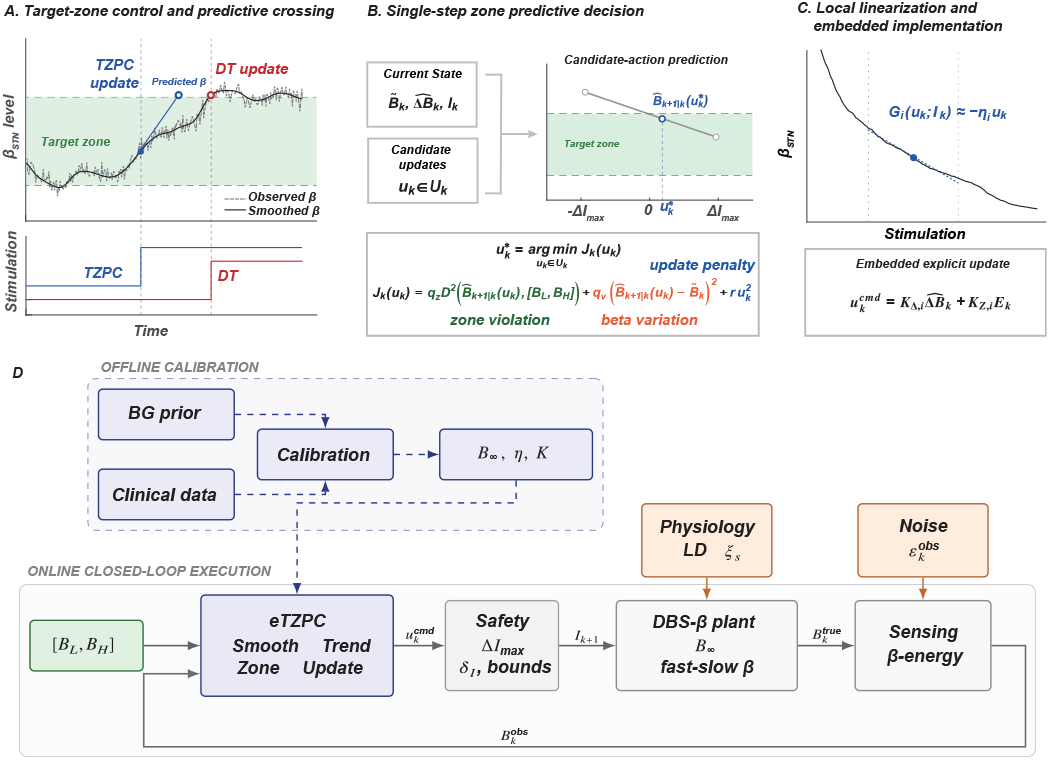
Trend–zone predictive control design and embedded implementation. (A) Predictive crossing response compared with DT control. (B) Single-step TZPC decision over candidate stimulation updates. (C) Local linearization yielding the explicit eTZPC law. (D) Offline calibration and online constrained beta-feedback execution. BG, basal ganglia; DBS, deep brain stimulation; DT, dual-threshold control; TZPC, trend– zone predictive control; eTZPC, embedded TZPC.

The controller predicts the *β*_STN_ level in the next control cycle for candidate stimulation updates, and selects the optimal action according to target-zone violation, predicted beta change, and stimulation-update cost (Fig. 2B). This structure directly incorporates the patient-specific stimulation– *β*_STN_ mapping into the control decision, such that stimulation updates are simultaneously constrained by the target zone, single-step amplitude limit, and safety boundaries.

To meet the online operation requirements of implantable devices, the full TZPC was further reduced to an explicit embedded control law (Fig. 2C). By approximating local stimulation sensitivity within the clinical stimulation working range of each patient, eTZPC simplified online computation to observation smoothing, trend estimation, zone-error calculation, fixed-gain updating, amplitude quantization, and boundary checking, without requiring online nonlinear model evaluation or candidate-action search.

### C. eTZPC Achieved a Better Trade-off Between Target-Zone Modulation and Stimulation-Switching Burden

To evaluate closed-loop control performance, we compared DT, PID, and eTZPC under the four test scenarios described above, using the same patient-specific target zones, stimulation safety boundaries, maximum single-step change constraint, stimulation-amplitude resolution, and one minute’s update interval. Representative trajectories showed that DT exhibited reactive, infrequent updates after threshold violation, PID produced more continuous and frequent amplitude adjustments, whereas eTZPC generated discrete constrained updates based on smoothing, trend prediction, and zone error (Fig. 3A).

**Fig. 3.**
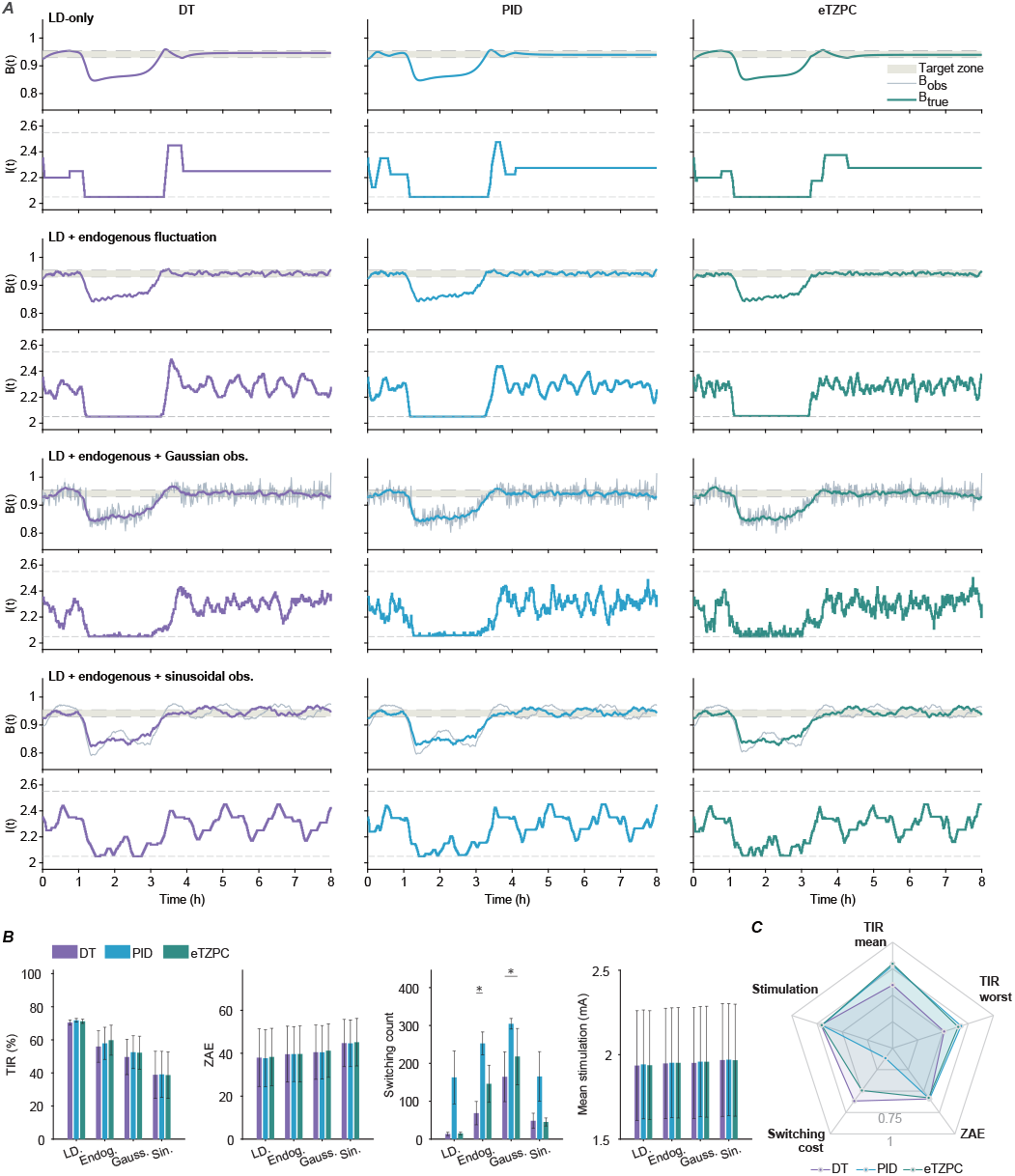
Closed-loop control performance across disturbance scenarios. (A)Representative 8-h beta and stimulation trajectories of DT, PID, and eTZPC under four disturbance scenarios. Gray traces indicate observed beta, colored traces indicate latent beta, and dashed lines indicate stimulation bounds. (B) Quantitative comparison of TIR, ZAE, switching count, and mean stimulation amplitude across controllers and scenarios. Bars show mean ± SD across patients; asterisks indicate significant pairwise differences. (C) Radar summary of normalized performance across key metrics. TIR, time-in-range ratio; ZAE, zone absolute error

The zone-control metrics showed that eTZPC achieved TIR values close to those of PID across the four scenarios, and outperformed DT in some disturbance scenarios (Fig. 3B). In the LD-only scenario, the TIR values of all three controllers were approximately 70%, suggesting that under low-disturbance conditions DT was already able to achieve basic target-zone maintenance. Under the endogenous-fluctuation and Gaussian-observation-noise scenarios, the TIR values of eTZPC were 59.8± 20.3% and 52.2 ±22.4%, respectively, which were higher than those of DT (56.0 ±21.2% and 49.7 ±23.9%) and close to those of PID (57.9 ± 21.7% and 52.5 ± 22.2%). In the sinusoidal observation disturbance scenario, the TIR values of all three controllers decreased to approximately 39%, indicating that strong periodic observation disturbance simultaneously limited the target-zone maintenance ability of all feedback strategies. ZAE did not consistently favor eTZPC over PID, but overall supported the interpretation that eTZPC achieved zone-control performance close to PID and comparable to DT.

Stimulation-switching burden further distinguished the controllers. Compared with PID, eTZPC reduced switching burden while maintaining a similar TIR (Fig. 3B). This difference was already evident in the LD-only scenario, where the switching counts of PID and eTZPC were 163.0 ± 156.5 and 15.0 ± 7.1, respectively; in the Gaussian observation noise scenario, they were 304.5 ± 31.2 and 218.1± 165.7, respectively; and in the sinusoidal observation disturbance scenario, they were 165.3 ± 146.9 and 45.3 ± 23.5, respectively. Meanwhile, in the Gaussian observation noise scenario, the in-zone update rate of eTZPC was lower than that of PID (0.226 ± 0.174 vs. 0.450 ± 0.223), suggesting that its smoothing, trend prediction, and constrained zone-feedback structure helped reduce noise-driven false stimulation triggering.

The radar plots further illustrated the trade-off advantage of eTZPC (Fig. 3C). Compared with DT, eTZPC achieved higher TIR in most disturbance scenarios; compared with PID, eTZPC maintained a lower switching burden. Overall, although eTZPC did not comprehensively outperform all controllers on every single metric, it achieved a more balanced profile, combining PID-like target-zone regulation with DT-like low-switching output.

### D. Progressive Ablation Analysis Showed That Each eTZPC Module Addressed a Distinct Closed-Loop Failure Mode

To clarify the role of each component of eTZPC, a progressive ablation analysis was performed (Fig. 4A–D).

**Fig. 4.**
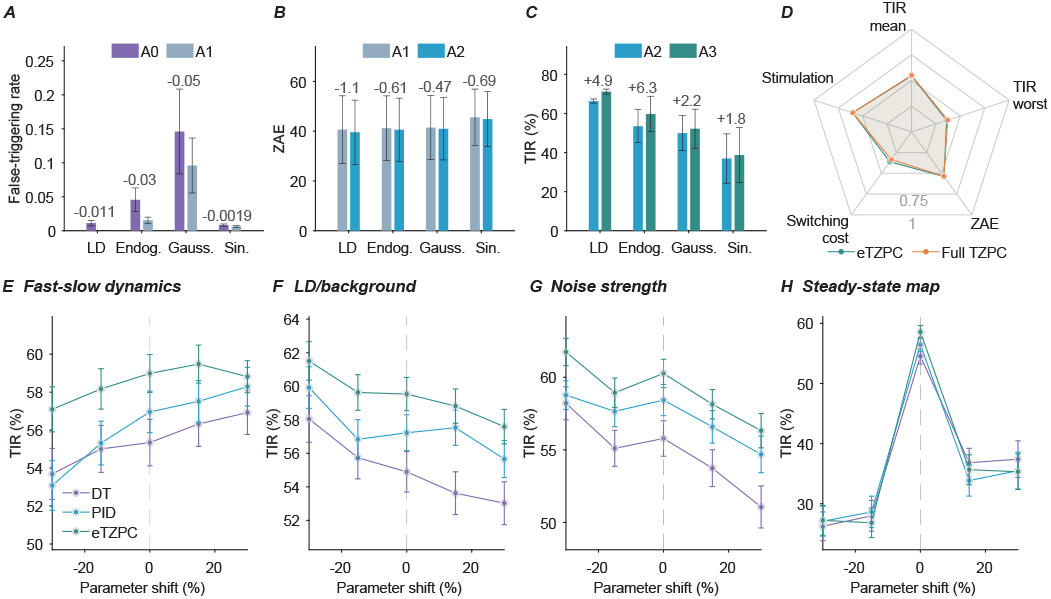
Component ablation and parameter-mismatch robustness of eTZPC. (A)–(D) Effects of observation smoothing, trend prediction, patient-specific action prediction, and embedded implementation. (E)–(H) TIR under mismatch in fast–slow dynamics, LD/background modulation, noise strength, and the steady-state stimulation–***β*** map. The vertical dashed line indicates the nominal calibrated model. Bars and points show mean **±** SEM.

Observation smoothing mainly reduced noise-driven false triggering and stimulation switching, but introduced feedback delay when used alone. To quantify this effect, we additionally calculated the false-trigger rate in the comparison between A0 and A1, defined as the proportion of stimulation updates triggered by *B*_obs_ while the latent state *B*_true_ remained within the target zone. In the Gaussian observation noise scenario, A1 reduced the switching burden from 164.7 to 103.4 and reduced the false-trigger rate from 0.146 to 0.096 compared with A0 (Fig. 4A), but did not improve TIR. After trend prediction was added, A2 further reduced the zone violation area, indicating that one-step no-action prediction partially compensated for the feedback delay introduced by observation smoothing (Fig. 4B).

Patient-specific action prediction produced the largest recovery in TIR. Compared with A2, the full TZPC formulation (A3) increased TIR by 4.9, 6.3, 2.2, and 1.8 percentage points across the four scenarios, respectively (Fig. 4C). This indicates that the patient-specific stimulation– *β*_STN_ response model is an important component for maintaining zone control under disturbance conditions, and that smoothing and trend prediction alone were insufficient to recover the full TZPC performance. Finally, the embedded explicit implementation (A4/eTZPC) largely preserved the overall control performance of A3. The radar plot showed that A4 and A3 were broadly comparable in TIR, zone error, switching burden, and stimulation level (Fig. 4D), suggesting that the full TZPC could be compressed into an embedded control law under local linearization and fixed-gain constraints without substantially degrading the main closed-loop behavior.

Overall, the ablation results showed that different modules of eTZPC addressed distinct closed-loop failure modes: observation smoothing reduced noise-driven false triggering, trend prediction alleviated feedback delay, patient-specific action prediction mitigated stimulation–*β*_STN_ response mismatch, and embedded explicit implementation supported constrained online execution while preserving the main control behavior.

### E. eTZPC Maintained Relatively Stable TIR Under Most Model-Mismatch Scenarios

To evaluate the effect of patient-model calibration errors on closed-loop control, we perturbed only the closed-loop plant parameters while keeping the controller parameters fixed. Four classes of mismatch were tested: fast–slow stimulation-response dynamics, LD/background modulation, noise strength, and stimulation–*β*_STN_ steady-state mapping (Fig. 4E–H). Under fast–slow dynamic mismatch, the TIR of eTZPC changed smoothly with parameter shifts and remained higher than those of DT and PID at most perturbation levels (Fig. 4E). Under LD/background modulation mismatch, eTZPC maintained relatively high TIR across most perturbation levels, suggesting a degree of tolerance to deviations in medication- and background-related slow modulation amplitude (Fig. 4F). Under noise-strength mismatch, eTZPC also maintained relatively stable TIR, whereas DT showed a more pronounced decline as noise increased (Fig. 4G). Steady-state mapping mismatch had a substantial effect on all three controllers, indicating that the patient-specific stimulation– *β*_STN_ steady-state map remains a key calibration target for closed-loop performance (Fig. 4H). Overall, eTZPC showed more gradual performance changes when fast–slow dynamics, medication/background modulation, and noise strength deviated from the calibrated values, but its performance still depended on reliable calibration of the patient-specific steady-state stimulation–*β*_STN_ map.

### F. Embedded Implementation Reduced Online Complexity and Preserved Execution Consistency in Patient-in-the-Loop Data

Finally, we evaluated whether the full TZPC could be reduced to an embedded explicit controller suitable for operation in an implantable device (Fig. 5A–D). The full TZPC requires evaluation of candidate stimulation actions and their predicted beta responses at each control step, whereas eTZPC reduces this process to an explicit trend–zone feedback law. During online operation, eTZPC only requires observation smoothing, trend estimation, zone-error calculation, stimulation updating, quantization, and boundary checking.

**Fig. 5.**
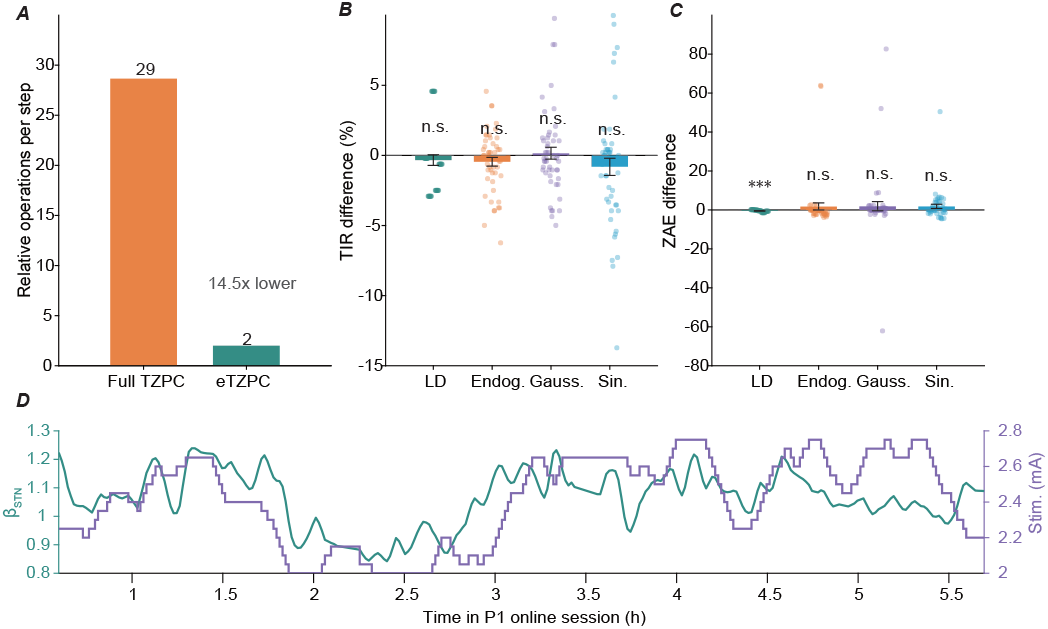
Simulation-level embedded-reduction analysis and on-device eTZPC execution consistency. (A) Simulation-level online-operation proxy for full TZPC and eTZPC, reduced from 29 to 2 relative operations per step. (B)–(C) Simulation-level differences in TIR and ZAE between eTZPC and full TZPC across disturbance scenarios. Dots indicate individual results; error bars indicate mean **±** SEM. (D) Representative patient-in-the-loop recording during on-device eTZPC operation, showing device-recorded ***β***_**STN**_ and delivered stimulation amplitude. Only eTZPC was implemented on the implantable pulse generator.

The computational-burden comparison showed that the relative online operation proxy of the full TZPC was 29 operations per step, whereas that of eTZPC was reduced to 2 operations per step, corresponding to an approximately 14.5-fold reduction (Fig. 5A). This result indicates that eTZPC avoids online candidate-action search and nonlinear action-effect computation, making it more suitable for minute-scale embedded cDBS implementation at the algorithmic level.

Further simulation comparison showed that the embedded explicit implementation largely preserved the main closed-loop behavior of the full TZPC (Fig. 5B–C). Across the four test scenarios, the TIR differences between eTZPC and the full TZPC did not reach statistical significance (Fig. 5B), suggesting that local linearization and fixed-gain implementation did not substantially impair target-zone maintenance. The ZAE difference was also not significant in most disturbance scenarios. A statistically significant difference was observed only in the LD-only scenario, but its absolute magnitude was small (Fig. 5C). Thus, the reduced-order implementation of eTZPC did not cause systematic degradation of the main closed-loop performance.

It should be emphasized that the comparisons in Fig. 5A–C were simulation-level analyses between full TZPC and its embedded explicit approximation, whereas only eTZPC was implemented on the implantable pulse generator. Therefore, Fig. 5B–C evaluate whether embedded reduction preserved the control behavior of full TZPC, while Fig. 5D evaluates on-device execution of the selected eTZPC algorithm rather than a hardware-level comparison among controllers.

Real patient-in-the-loop validation further confirmed embedded feasibility (Fig. 5D). eTZPC completed *>*8-h online closed-loop operation in all five patients, with delivered stimulation amplitudes satisfying predefined safety boundaries and maximum single-step change constraints. Offline replay of device-recorded biomarkers showed 100% stimulation-update consistency within a continuous valid segment and 97.29% consistency across the full-day recording. The remaining discrepancies were attributable to missing recorded data rather than errors in the eTZPC decision rule.

## IV. Discussion

Bridging mechanistic neural models with the deployment constraints of implantable DBS devices remains a central challenge for closed-loop stimulation. In this study, we aimed to address this gap by deriving an embedded explicit trend-zone predictive controller, eTZPC, based on a patient-calibrated DBS–*β*_STN_ dynamical model. In addition, we compared eTZPC with representative control strategies, including DT and PID control, in terms of robustness, constraint satisfaction, and engineering deployability, demonstrating its advantages in reducing stimulation-switching burden and maintaining robust target-zone modulation.

The basal-ganglia computational model was used here as a mechanistic prior for the stimulation–*β*_STN_ steady-state relationship, and after alignment with patient stimulation–response recordings, this prior provided individualized steady-state maps that were used to define the target zone, local stimulation sensitivity, closed-loop simulation plant, and parameter-mismatch tests. This design constrained calibration from sparse clinical amplitude-response points and reduced reliance on an unconstrained local linear fit.

Within this framework, eTZPC was introduced. eTZPC can be interpreted as the projection of a patient-specific predictive control problem onto the class of feedback laws executable on the implant, thereby distinguishing it from a “smoothed DT rule” or a “simplified MPC”. The full TZPC jointly accounts for predicted zone violation, expected biomarker change, update cost, and embedded hardware constraints when selecting stimulation updates. eTZPC then uses local linearization to convert the same objective into a fixed-gain, quantized, safety-constrained feedback law. Crucially, individualization is preserved at a level that DT control cannot reach: compared with DT’s boundary-only individualization, eTZPC further enables patient-specific stimulation-update responses through the calibrated stimulation–*β*_STN_ map. Thus, eTZPC differs from DT control in principle by incorporating patient-specific stimulation-response sensitivity into the update law.

In real-world cDBS deployment, controller performance should be assessed beyond any single metric. A clinically relevant controller needs to balance target-zone regulation with device-level constraints and patient tolerance. The comparison among DT, PID, and eTZPC is best read in this light: as a deployment-constrained trade-off, without implying universal superiority. eTZPC achieved target-zone regulation comparable to PID in most scenarios, while reducing stimulation-switching burden relative to PID and preserving constraint satisfaction. This distinction is clinically relevant because frequent amplitude updates may increase patient intolerance and device-operation burden even when each individual update remains within the nominal safety range. Therefore, we adopted a 1-min update interval, a commonly used timescale in clinical cDBS that balances device capability and patient tolerability. Compared with DT control, eTZPC added trend-based and patient-specific action components, which improved performance in several disturbed conditions but did not eliminate the effect of strong observation disturbance. Parameter-mismatch tests further indicated that reliable calibration of the steady-state stimulation–*β*_STN_ map remained important, whereas deviations in fast–slow dynamics, medication/background modulation, and noise strength produced more gradual performance changes. This profile is itself a deployment-favorable trade-off: eTZPC concentrates its calibration demand on the single component that is most readily obtained in clinical practice, a one-time amplitude sweep, and tolerates approximation in precisely those components that are hardest to identify from chronic recordings.

This study has several limitations. First, the model incorporated multiscale features to a certain extent through the steady-state stimulation–*β*_STN_ map and fast–slow responses after stimulation changes, but it was still applied mainly to minute-scale stimulation control and relied on lightweight smoothing and local trend estimation. Future work will consider longer-timescale endogenous dynamical changes and further developing larger-scale adaptive modeling and control frameworks, including adaptive parameter updating, richer state representations, and multi-objective control [32], [33]. Second, eTZPC should be viewed as a proof-of-concept embedded controller, and the sample size used for evaluation remains limited. Larger longitudinal studies are required to validate its generalizability and clinical benefit. Third, although cDBS ultimately aims to adjust stimulation according to the patient’s neural and clinical state [34], *β*_STN_ was used here as a simplified biomarker-level proxy. Because biomarker dynamics may not be temporally aligned with state changes, target-zone *β*_STN_ regulation cannot fully represent state control or be directly equated with clinical benefit. This limitation may be further amplified by intersubject variability in *β*_STN_-band responsiveness, stimulation sensitivity, and levodopa-related pharmacokinetic or pharmacodynamic profiles.

## V. Conclusion

This study developed a patient-calibrated *β*_STN_ dynamical model and an embedded explicit trend–zone predictive controller for closed-loop DBS. eTZPC achieved a deployment-oriented trade-off between PID-like target-zone beta regulation and reduced stimulation-switching burden, showed tolerance to dynamic and disturbance-parameter mismatch, and satisfied stimulation-boundary and step-size constraints during patient-in-the-loop recordings. These findings support patient-calibrated dynamical modeling combined with low-complexity explicit control as a feasible framework for embedded minute-scale cDBS evaluation.

## Acknowledgment

The authors would like to thank Prof. Jianguo Zhang and Prof. Fangang Meng from Beijing Tiantan Hospital for their support in surgical implantation and patient management.

The authors also thank Haiyu Tian, Yingtian Liu, Guokun Zhang from the National Engineering Research Center for Neuromodulation (NERCN) for their assistance in data acquisition.

## Notes

This work was supported in part by the National Key Research and Development Program of China under Grant 2022YFC2405100. This study involved human participants. Approval of all ethical and experimental procedures and protocols was granted by the Ethics Committee of Beijing Tiantan Hospital (Approval No. KY2023-158-02), and conducted in accordance with the Declaration of Helsinki.

### Competing Interest Statement

The authors have declared no competing interest.

