## Supplementary File for "Patient-Calibrated Dynamical Modeling and Embedded Trend-Zone Predictive Control for Closed-Loop Deep Brain Stimulation in Parkinson’s Disease"

**Supplementary Methods**

S1 Basal ganglia network model used as a mechanistic prior

The basal ganglia network model was adapted from the Rubin–Terman model of pathological basal-ganglia–thalamocortical activity.

The spiking basal ganglia model was used to generate a mechanistically constrained prior for the steady-state relationship between stimulation amplitude and simulated STN beta power. The model was not used as the online plant in closed-loop simulations and was not embedded in the controller. Instead, its averaged stimulation–beta curve was subsequently aligned to patient recordings to obtain patient-specific steady-state maps.

S1.1 Neural population equations and parameters

The membrane potential dynamics of STN neurons were modeled as:

$$C_{m}\frac{dv}{dt}=-I_{L}-I_{K}-I_{Na}-I_{T}-I_{Ca}-I_{AHP}-I_{syn_{G\to S}}+I_{app-STN}+I_{input}.$$

The ionic-current equations and parameter values are summarized in Supplementary Tables S1 and S2.

TABLE S1 Current Equation of STN Neurons

| **Current Equations** | **Gating Variable Equations** | | | |
| --- | --- | --- | --- | --- |
| $I_{L}=g_{L}\left( v-V_{L} \right)$ |  | |  | |
| $I_{K}=g_{K}n^{4}\left( v-V_{k} \right)$ | $\frac{dn}{dt}=\phi_{n}\frac{n_{\infty}-n}{\tau_{n}}$ | $\tau_{n}=\tau_{n}^{0}+\frac{\tau_{n}^{1}}{1+e^{-\frac{v-\theta_{n}^{\tau}}{\sigma_{n}^{\tau}}}}$ | | $n_{\infty}=\frac{1}{1+e^{-\frac{v-\theta_{n}}{\sigma_{n}}}}$ |
| $I_{Na}=g_{Na}m_{\infty}^{3}h\left( v-V_{Na} \right)$ | $\frac{dh}{dt}=\phi_{h}\frac{h_{\infty}-h}{\tau_{h}}$ | $\tau_{h}=\tau_{h}^{0}+\frac{\tau_{h}^{1}}{1+e^{-\frac{v-\theta_{h}^{\tau}}{\sigma_{h}^{\tau}}}}$ | | $h_{\infty}=\frac{1}{1+e^{-\frac{v-\theta_{h}}{\sigma_{h}}}}$ |
|  | $m_{\infty}=\frac{1}{1+e^{-\frac{v-\theta_{m}}{\sigma_{m}}}}$ |  | |  |
| $I_{T}=g_{T}a_{\infty}^{3}b_{\infty}^{2}(v-V_{Ca})$ | $a_{\infty}=\frac{1}{1+e^{-\frac{v-\theta_{a}}{\sigma_{a}}}}$ | $b_{\infty}=\frac{1}{1+e^{-\frac{r-\theta_{b}}{\sigma_{b}}}}-\frac{1}{1+e^{-\frac{\theta_{b}}{\sigma_{b}}}}$ | | |
|  | $\frac{dr}{dt}=\phi_{r}\frac{r_{\infty}-r}{\tau_{r}}$ | $\tau_{r}=\tau_{r}^{0}+\frac{\tau_{r}^{1}}{1+e^{-\frac{v-\theta_{r}^{\tau}}{\sigma_{r}^{\tau}}}}$ | | $r_{\infty}=\frac{1}{1+e^{-\frac{v-\theta_{r}}{\sigma_{r}}}}$ |
| $I_{Ca}=g_{Ca}s_{\infty}^{2}\left( v-V_{Ca} \right)$ | $s_{\infty}=\frac{1}{1+e^{-\frac{v-\theta_{s}}{\sigma_{s}}}}$ | | | |
| $I_{AHP}=g_{AHP}\left( v-V_{K} \right)\cdot\frac{\left[ Ca \right]}{\left[ Ca \right]+k_{1}}$ | $\frac{d\left[ Ca \right]}{dt}=\epsilon\left( -I_{Ca}-I_{T}-k_{Ca}\left[ Ca \right] \right)$ | | | |

TABLE S2 Parameter values of STN Neurons

| **Parameter** | | **Value** | | **Parameter** | | **Value** | | **Parameter** | | **Value** | | **Parameter** | | **Value** | |
| --- | --- | --- | --- | --- | --- | --- | --- | --- | --- | --- | --- | --- | --- | --- | --- |
| $C_{m}$ | 1 | | $pF/\mu m^{2}$ | $\tau_{n}^{0}$ | 1.0 | | $ms$ | $\theta_{n}$ | -32.0 | | $mV$ | $\sigma_{n}$ | 8.0 | | $mV$ |
| $g_{L}$ | 2.25 | | $nS/\mu m^{2}$ | $\tau_{h}^{0}$ | 1.0 | | $ms$ | $\theta_{h}$ | -39.0 | | $mV$ | $\sigma_{h}$ | -3.1 | | $mV$ |
| $g_{K}$ | 45.0 | | $nS/\mu m^{2}$ | $\tau_{r}^{0}$ | 40.0 | | $ms$ | $\theta_{r}$ | -67.0 | | $mV$ | $\sigma_{r}$ | -2.0 | | $mV$ |
| $g_{Na}$ | 37.5 | | $nS/\mu m^{2}$ | $\tau_{n}^{1}$ | 100.0 | | $ms$ | $\theta_{a}$ | -63.0 | | $mV$ | $\sigma_{a}$ | 7.8 | | $mV$ |
| $g_{T}$ | 0.5 | | $nS/\mu m^{2}$ | $\tau_{h}^{1}$ | 500.0 | | $ms$ | $\theta_{m}$ | -30.0 | | $mV$ | $\sigma_{m}$ | 15.0 | | $mV$ |
| $g_{Ca}$ | 0.5 | | $nS/\mu m^{2}$ | $\tau_{r}^{1}$ | 17.5 | | $ms$ | $\theta_{s}$ | -39.0 | | $mV$ | $\sigma_{s}$ | 8.0 | | $mV$ |
| $g_{AHP}$ | 9.0 | | $nS/\mu m^{2}$ | $\phi_{n}$ | 0.75 | |  | $\theta_{b}$ | 0.4 | | $mV$ | $\sigma_{b}$ | -0.1 | | $mV$ |
| $I_{app-STN}$ | 25.0 | | $pA/\mu m^{2}$ | $\phi_{h}$ | 0.75 | |  | $\theta_{n}^{\tau}$ | -80.0 | | $mV$ | $\sigma_{n}^{\tau}$ | -26.0 | | $mV$ |
| $k_{1}$ | 15.0 | | $pA/\mu m^{2}$ | $\phi_{r}$ | 0.2 | |  | $\theta_{h}^{\tau}$ | -57.0 | | $mV$ | $\sigma_{h}^{\tau}$ | -3.0 | | $mV$ |
| $V_{Na}$ | 55.0 | | $mV$ | $V_{L}$ | -60.0 | | $mV$ | $\theta_{r}^{\tau}$ | 68.0 | | $mV$ | $\sigma_{r}^{\tau}$ | -2.2 | | $mV$ |
| $V_{Ca}$ | 140.0 | | $mV$ | $V_{k}$ | -80.0 | | $mV$ | $k_{Ca}$ | 22.5 | |  | $\epsilon$ | 3.75 | | $ms^{-1}$ |

The membrane potential dynamics of GPe neurons were described as:

$$C_{m}\frac{dv}{dt}=-I_{L}-I_{K}-I_{Na}-I_{T}-I_{Ca}-I_{AHP}-I_{\mathrm{syn}(STN\to GPe)}-I_{\mathrm{syn}(GPe\to GPe)}+I_{\mathrm{app},\mathrm{GPe}}.$$

GPi neurons were modeled using the same ionic-current structure as GPe neurons, but with nucleus-specific synaptic inputs and applied current. Specifically, $I_{\mathrm{syn}(STN\to GPe)}$, $I_{\mathrm{syn}(GPe\to GPe)}$, and $I_{\mathrm{app},\mathrm{GPe}}$were replaced by $I_{\mathrm{syn}(STN\to GPi)}$, $I_{\mathrm{syn}(GPe\to GPi)}$, and $I_{\mathrm{app},\mathrm{GPi}}$, respectively. The ionic-current equations and parameter values for GP neurons are summarized in Supplementary Tables S3 and S4.

TABLE S3 Current Equation of GP Neurons

| **Current Equations** | **Gating Variable Equations** | | | |
| --- | --- | --- | --- | --- |
| $I_{L}=g_{L}\left( v-V_{L} \right)$ |  | |  | |
| $I_{K}=g_{K}n^{4}\left( v-V_{k} \right)$ | $\frac{dn}{dt}=\phi_{n}\frac{n_{\infty}-n}{\tau_{n}}$ | $\tau_{n}=\tau_{n}^{0}+\frac{\tau_{n}^{1}}{1+e^{-\frac{v-\theta_{n}^{\tau}}{\sigma_{n}^{\tau}}}}$ | | $n_{\infty}=\frac{1}{1+e^{-\frac{v-\theta_{n}}{\sigma_{n}}}}$ |
| $I_{Na}=g_{Na}m_{\infty}^{3}h\left( v-V_{Na} \right)$ | $\frac{dh}{dt}=\phi_{h}\frac{h_{\infty}-h}{\tau_{h}}$ | $\tau_{h}=\tau_{h}^{0}+\frac{\tau_{h}^{1}}{1+e^{-\frac{v-\theta_{h}^{\tau}}{\sigma_{h}^{\tau}}}}$ | | $h_{\infty}=\frac{1}{1+e^{-\frac{v-\theta_{h}}{\sigma_{h}}}}$ |
|  | $m_{\infty}=\frac{1}{1+e^{-\frac{v-\theta_{m}}{\sigma_{m}}}}$ |  | |  |
| $I_{T}=g_{T}a_{\infty}^{3}r(v-V_{Ca})$ | $\frac{dr}{dt}=\phi_{r}\frac{r_{\infty}-r}{\tau_{r}}$ | $r_{\infty}=\frac{1}{1+e^{-\frac{v-\theta_{r}}{\sigma_{r}}}}$ | | $a_{\infty}=\frac{1}{1+e^{-\frac{v-\theta_{a}}{\sigma_{a}}}}$ |
| $I_{Ca}=g_{Ca}s_{\infty}^{2}\left( v-V_{Ca} \right)$ | $s_{\infty}=\frac{1}{1+e^{-\frac{v-\theta_{s}}{\sigma_{s}}}}$ | | | |
| $I_{AHP}=g_{AHP}\left( v-V_{K} \right)\cdot\frac{\left[ Ca \right]}{\left[ Ca \right]+k_{1}}$ | $\frac{d\left[ Ca \right]}{dt}=\epsilon\left( -I_{Ca}-I_{T}-k_{Ca}\left[ Ca \right] \right)$ | | | |

TABLE S4 Parameter values of GP Neurons

| **Parameter** | | **Value** | | **Parameter** | | **Value** | | **Parameter** | | **Value** | | **Parameter** | | | **Value** | |
| --- | --- | --- | --- | --- | --- | --- | --- | --- | --- | --- | --- | --- | --- | --- | --- | --- |
| $C_{m}$ | 1 | | $pF/\mu m^{2}$ | $\tau_{n}^{0}$ | 0.05 | | $ms$ | $\theta_{n}$ | -50.0 | | $mV$ | $\sigma_{n}$ | 14.0 | | | $mV$ |
| $g_{L}$ | 0.1 | | $nS/\mu m^{2}$ | $\tau_{h}^{0}$ | 0.05 | | $ms$ | $\theta_{h}$ | -58.0 | | $mV$ | $\sigma_{h}$ | -12.0 | | | $mV$ |
| $g_{K}$ | 30.0 | | $nS/\mu m^{2}$ | $\tau_{n}^{1}$ | 0.27 | | $ms$ | $\theta_{r}$ | -70.0 | | $mV$ | $\sigma_{r}$ | -2.0 | | | $mV$ |
| $g_{Na}$ | 120.0 | | $nS/\mu m^{2}$ | $\tau_{h}^{1}$ | 0.27 | | $ms$ | $\theta_{a}$ | -57.0 | | $mV$ | $\sigma_{a}$ | 2.0 | | | $mV$ |
| $g_{T}$ | 0.5 | | $nS/\mu m^{2}$ | $V_{Na}$ | 55.0 | | $mV$ | $\theta_{m}$ | -37.0 | | $mV$ | $\sigma_{m}$ | 10.0 | | | $mV$ |
| $g_{Ca}$ | 0.15 | | $nS/\mu m^{2}$ | $V_{Ca}$ | 120.0 | | $mV$ | $\theta_{s}$ | -35.0 | | $mV$ | $\sigma_{s}$ | 2.0 | | | $mV$ |
| $g_{AHP}$ | 30.0 | | $nS/\mu m^{2}$ | $V_{L}$ | -55.0 | | $mV$ | $\theta_{n}^{\tau}$ | -40.0 | | $mV$ | $\sigma_{n}^{\tau}$ | | -12.0 | | $mV$ |
| $I_{app-GPe}$ | 2.0 | | $pA/\mu m^{2}$ | $V_{k}$ | -80.0 | | $mV$ | $\theta_{h}^{\tau}$ | -40.0 | | $mV$ | $\sigma_{h}^{\tau}$ | -12.0 | | | $mV$ |
| $I_{app-GPi}$ | 3.0 | | $pA/\mu m^{2}$ | $k_{Ca}$ | 15.0 | |  | $\phi_{n}$ | 0.1 | |  | $\phi_{r}$ | 1.0 | | |  |
| $k_{1}$ | 30.0 | | $pA/\mu m^{2}$ | $\epsilon$ | 0.0001 | | $ms^{-1}$ | $\phi_{h}$ | 0.05 | |  |  |  | | |  |

The membrane potential dynamics of thalamic neurons were described as:

$$C_{m}\frac{dv}{dt}=-I_{L}-I_{K}-I_{Na}-I_{T}-I_{\mathrm{syn}(GPi\to Th)}+I_{\mathrm{SM}}.$$

Here, $I_{\mathrm{SM}}$denotes the sensorimotor cortical input to thalamic neurons. The ionic-current equations and parameter values for thalamic neurons are summarized in Supplementary TABLEs S5 and S6.

TABLE S5 Current Equation of TH Neurons

| **Current Equations** | **Gating Variable Equations** | | | |
| --- | --- | --- | --- | --- |
| $I_{L}=g_{L}\left( v-V_{L} \right)$ |  | |  | |
| $I_{K}=g_{K}\left( 0.75\left( 1-h \right) \right)^{4}\left( v-V_{k} \right)$ | $\frac{dh}{dt}=\frac{h_{\infty}-h}{\tau_{h}}$ | $h_{\infty}=\frac{1}{1+e^{-\frac{v-\theta_{h}}{\sigma_{h}}}}$ | | $\tau_{h}=\frac{1}{a_{h}+b_{h}}$ |
|  |  | $a_{h}=\tau_{h}^{a}e^{-\frac{v-\theta_{a}^{\tau}}{\sigma_{a}^{\tau}}}$ | | $b_{h}=\frac{\tau_{h}^{b}}{1+e^{-\frac{v-\theta_{b}^{\tau}}{\sigma_{b}^{\tau}}}}$ |
| $I_{Na}=g_{Na}m_{\infty}^{3}h\left( v-V_{Na} \right)$ | $m_{\infty}=\frac{1}{1+e^{-\frac{v-\theta_{m}}{\sigma_{m}}}}$ |  | |  |
| $I_{T}=g_{T}p_{\infty}^{3}r(v-V_{Ca})$ | $\frac{dr}{dt}=\frac{r_{\infty}-r}{\tau_{r}}$ | $\tau_{r}=\tau_{r}^{0}+e^{-\frac{v-\theta_{r}^{\tau}}{\sigma_{r}^{\tau}}}$ | | $r_{\infty}=\frac{1}{1+e^{-\frac{v-\theta_{r}}{\sigma_{r}}}}$ |
|  | $p_{\infty}=\frac{1}{1+e^{-\frac{v-\theta_{p}}{\sigma_{p}}}}$ |  | |  |
| $I_{SM}=i_{SM}H(sin(\frac{2\pi t}{\rho_{SM}}))\times[1 - H(sin(\frac{2\pi\left( t + \delta_{SM} \right)}{\rho_{SM}}))$ | | | | |

TABLE S6 Parameter values of TH Neurons

| **Parameter** | | **Value** | | **Parameter** | | **Value** | | **Parameter** | | **Value** | | **Parameter** | | **Value** | |
| --- | --- | --- | --- | --- | --- | --- | --- | --- | --- | --- | --- | --- | --- | --- | --- |
| $C_{m}$ | 1 | | $pF/\mu m^{2}$ | $\tau_{r}^{0}$ | 28.0 | | $ms$ | $\theta_{r}$ | -84.0 | | $mV$ | $\sigma_{r}$ | -4.0 | | $mV$ |
| $g_{L}$ | 0.05 | | $nS/\mu m^{2}$ | $\tau_{h}^{a}$ | 0.128 | | $ms^{-1}$ | $\theta_{r}^{\tau}$ | -25.0 | | $mV$ | $\sigma_{r}^{\tau}$ | 10.5 | | $mV$ |
| $g_{K}$ | 5.0 | | $nS/\mu m^{2}$ | $\tau_{h}^{b}$ | 4.0 | | $ms^{-1}$ | $\theta_{a}^{\tau}$ | -46.0 | | $mV$ | $\sigma_{a}^{\tau}$ | 18.0 | | $mV$ |
| $g_{Na}$ | 3.0 | | $nS/\mu m^{2}$ | $V_{Na}$ | 50.0 | | $mV$ | $\theta_{b}^{\tau}$ | -23.0 | | $mV$ | $\sigma_{b}^{\tau}$ | 5.0 | | $mV$ |
| $g_{T}$ | 5.0 | | $nS/\mu m^{2}$ | $V_{T}$ | 0.0 | | $mV$ | $\theta_{p}$ | -60.0 | | $mV$ | $\sigma_{p}$ | 6.2 | | $mV$ |
| $I_{app-Th}$ |  | | $pA/\mu m^{2}$ | $V_{L}$ | -70.0 | | $mV$ | $\theta_{h}$ | -41.0 | | $mV$ | $\sigma_{h}$ | -4.0 | | $mV$ |
|  |  | |  | $V_{k}$ | -90.0 | | $mV$ | $\theta_{m}$ | -37.0 | | $mV$ | $\sigma_{m}$ | 7.0 | | $mV$ |

The synaptic current from population $\alpha$to population $\beta$was modeled as:

$$I_{\mathrm{syn}(\alpha\to\beta)}=g_{\alpha\to\beta}\left( v_{\alpha} - V_{\alpha\to\beta} \right)\sum_{j} s_{\alpha}^{j},$$

where $g_{\alpha\to\beta}$is the synaptic conductance, $V_{\alpha\to\beta}$is the reversal potential, and $s_{\alpha}^{j}$is the synaptic gating variable of the $j$-th presynaptic neuron in population $\alpha$.

The gating variable evolved according to:

$$\frac{ds_{\alpha}}{dt}=A_{\alpha}(1-s_{\alpha})H_{\infty}(v_{\alpha})-B_{\alpha}s_{\alpha}.$$

The steady-state activation function was defined as a sigmoid function:

$$H_{\infty}(v_{\alpha})=\frac{1}{1+\exp\left[ -(v_{\alpha}-\theta_{\alpha}-\theta_{\alpha}^{H})/\sigma_{\alpha}^{H} \right]}.$$

The parameters used for the synaptic-current equations are listed in Supplementary Table S7.

TABLE S7 Parameter values of synaptic functions

| **Parameter** | | **Value** | | **Parameter** | | **Value** | | **Parameter** | | **Value** | | **Parameter** | | **Value** |
| --- | --- | --- | --- | --- | --- | --- | --- | --- | --- | --- | --- | --- | --- | --- |
| $g_{GPe\to STN}$ | 0.9 | | $nS/\mu m^{2}$ | $V_{GPe\to STN}$ | -100.0 | | $mV$ | $\theta_{STN}$ | 30.0 | | $mV$ | $A_{STN}$ | 5.0 | |
| $g_{STN\to GPe}$ | 0.3 | | $nS/\mu m^{2}$ | $V_{STN\to GPe}$ | 0.0 | | $mV$ | $\theta_{GPe}$ | 20.0 | | $mV$ | $A_{GPe}$ | 2.0 | |
| $g_{GPe\to GPe}$ | 1.0 | | $nS/\mu m^{2}$ | $V_{GPe\to GPe}$ | -80.0 | | $mV$ | $\theta_{GPi}$ | 20.0 | | $mV$ | $A_{GPi}$ | 2.0 | |
| $g_{STN\to GPi}$ | 0.3 | | $nS/\mu m^{2}$ | $V_{STN\to GPi}$ | 0.0 | | $mV$ | $\theta_{STN}^{H}$ | -39.0 | | $mV$ | $B_{STN}$ | 1.0 | |
| $g_{GPe\to GPi}$ | 1.0 | | $nS/\mu m^{2}$ | $V_{GPe\to GPi}$ | -100.0 | | $mV$ | $\theta_{GPe,GPi}^{H}$ | -57.0 | | $mV$ | $B_{GPe}$ | 0.04 | |
| $g_{GPi\to Th}$ | 0.06 | | $nS/\mu m^{2}$ | $V_{GPi\to Th}$ | -85.0 | | $mV$ | $\sigma_{STN}^{H}$ | 8.0 | | $mV$ | $B_{GPi}$ | 0.08 | |
|  |  | |  |  |  | |  | $\sigma_{GPe,GPi}^{H}$ | 20.0 | | $mV$ |  |  | |

S1.2 Basal Ganglia Network Construction

The reduced basal ganglia network was constructed to preserve the relative composition of the four modeled nuclei while keeping the simulations computationally tractable. The numbers of neurons assigned to each nucleus are summarized in Supplementary Table S8.

TABLE S8 Neuron numbers in the basal ganglia network

| **STN** | **GPi** | **GPe** | **TH** |
| --- | --- | --- | --- |
| 28 | 16 | 72 | 14 |

For each simulation trial, synaptic connections among nuclei were randomly generated according to predefined projection rules. Each STN neuron received inhibitory input from two GPe neurons. Each GPe neuron received excitatory input from three STN neurons and inhibitory input from two other GPe neurons. Each GPi neuron received excitatory input from one STN neuron and inhibitory input from two GPe neurons. Each thalamic neuron received inhibitory input from two GPi neurons. Across 100 simulation trials, the average connection statistics are shown in Supplementary Fig. S1.

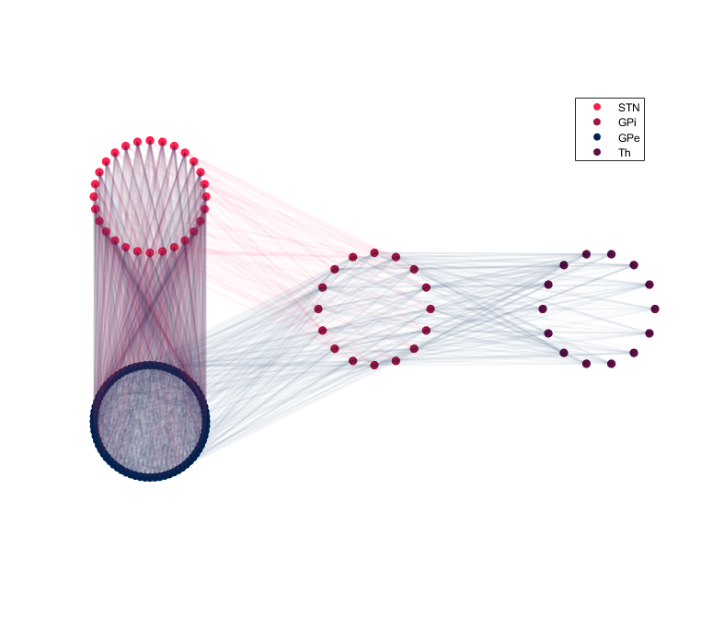

**Fig. S1. Basal ganglia network connectivity.** Connection statistics were averaged across 100 simulation trials with independently sampled synaptic connections.

S1.3 Electrode Geometry and STN Neuron Positioning

The inter-contact distance of the DBS lead was set to 2.25 mm. In each simulation trial, STN neurons were randomly distributed within a simplified rectangular volume of $9\times6\times4 \mathrm{mm}^{3}$centered on the stimulating contact C2. This geometry was used as an idealized spatial approximation of the STN region rather than a patient-specific anatomical reconstruction.

The stimulating contact C2 was placed at $\left( 0 , 0 , 0 \right)$mm. The two adjacent contacts used for bipolar LFP approximation were placed at $\left( -1.7559,-1.1706,-0.7804 \right)$mm for C1 and $\left( 1.7559 , 1.1706 , 0.7804 \right)$mm for C3. Neuron positions were independently resampled across simulation trials to account for spatial variability. The resulting average spatial distribution of simulated STN neurons is shown in Supplementary Fig. S2.

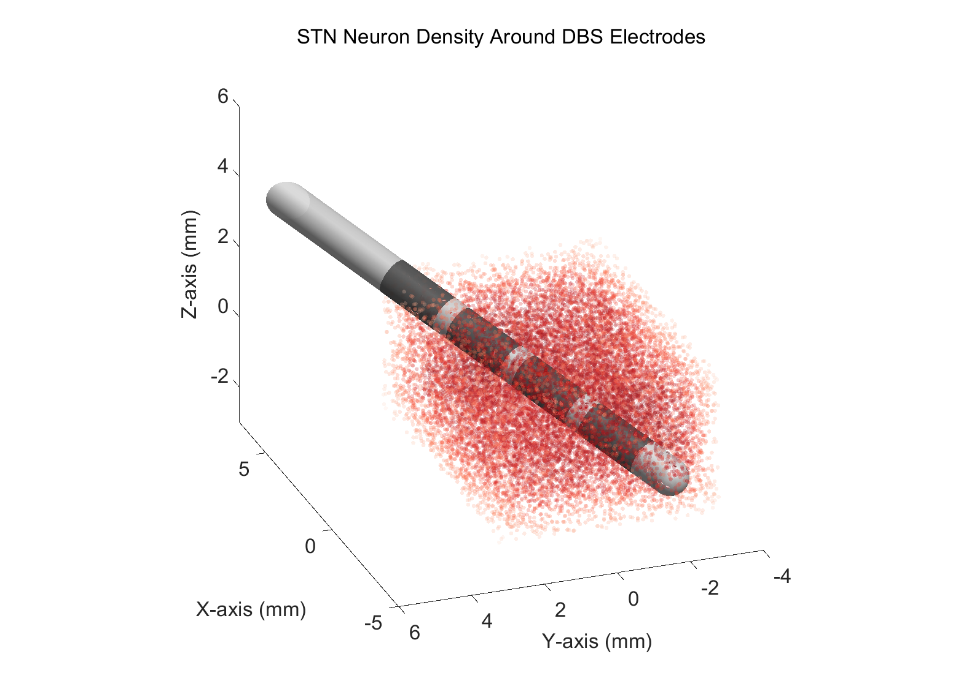

**Fig. S2. Simulated STN neuron density around DBS electrode contacts.** STN neurons were randomly distributed within an idealized rectangular STN volume across simulation trials.

S1.4 Generation of the Steady-State Stimulation–Beta Prior

For each stimulation amplitude, 100 independent simulation trials were performed with randomized synaptic connections and STN neuron positions. The simulation time step was set to 0.01 ms, corresponding to a sampling frequency of 100 kHz, and the total duration of each trial was 3000 ms. Membrane dynamics were integrated using the Euler method.

For each trial, the simulated bipolar local field potential (LFP) was obtained from the differential signal between the two recording contacts. Beta-band power was computed from the simulated LFP using Welch’s method. A 1-s segment length with 25% overlap was used for spectral estimation, and beta power was calculated by summing the power spectral density within the 13–30 Hz band.

For each stimulation amplitude, beta power was averaged across 100 independent trials to obtain a robust estimate of the steady-state stimulation–beta relationship. The resulting curve was smoothed and used as the computational prior $B_{\infty}^{\mathrm{prior}}(I)$for subsequent patient-specific calibration. This prior was used only to constrain the shape of the steady-state stimulation–beta map and was not used as the online closed-loop plant. In the volume-conduction approximation for LFP calculation, the tissue conductivity $\sigma$was set to $0.27 mS/mm$.

S1.5 Software and Tools for Basal Ganglia Modeling

The basal ganglia network simulations used to generate the steady-state stimulation–beta prior were implemented in Python 3.8.15 on a Linux platform. Neural network simulations were conducted using Brian2 2.4.2 with the cpp_standalone mode to improve computational efficiency. Numerical operations were performed using NumPy 1.23.5. Spectral estimation of simulated LFP signals was performed using the scipy.signal.welch function from SciPy 1.9.3.

S2 Clinical recordings and acquisition protocols

S2.1 Participants and Clinical Recordings

STN LFP recordings were obtained from participants with Parkinson’s disease implanted with bilateral STN DBS electrodes connected to a sensing-enabled implantable pulse generator (G106RS, Beijing PINS Medical Co., China). LFP signals were recorded directly through the implanted sensing system during stimulation-amplitude modulation and medication/background recording sessions.

S2.2 Stimulation-amplitude modulation recordings

For stimulation–beta calibration, recordings were performed in the MED OFF state to reduce confounding effects from dopaminergic medication. During each stimulation-amplitude modulation session, stimulation amplitude was adjusted stepwise within the clinically tolerated range, while stimulation contact configuration, frequency, and pulse width were kept unchanged. After each amplitude change, a 10-min wash-in period was recorded to characterize the transient beta response to stimulation adjustment. This wash-in segment was used to estimate the fast jump component and the minute-scale relaxation parameter $k$. After the wash-in period, an additional 5-min recording was obtained at the same stimulation amplitude and used to estimate the steady-state beta level for that amplitude. These steady-state beta estimates across stimulation amplitudes were then used to calibrate the patient-specific steady-state stimulation–beta map $B_{\infty}^{\left( i \right)}(I)$.

S2.3 Medication-cycle recordings

For medication-cycle recordings, participants withheld dopaminergic medication for at least 12 h before recording. Continuous STN LFP signals were then acquired from 30 min before medication intake to 120 min after medication intake, while stimulation parameters were kept unchanged. The 30-min pre-medication segment was used as the MED OFF baseline, and the post-medication recording window was used to characterize levodopa-related beta modulation over the onset, peak-effect, and washout phases. These recordings were used to estimate the patient-specific medication/background modulation term in the minute-scale beta model.

S2.4 Embedded closed-loop validation recordings

For embedded patient-in-the-loop closed-loop validation, the proposed controller was implemented in the implantable pulse generator and tested in all included patients during real online closed-loop operation. Each validation session lasted for more than 8 h. Patients withheld dopaminergic medication for at least 12 h before the start of the session, and then followed their normal daytime medication schedule during the closed-loop recording period. Thus, the online validation covered the transition from a MED OFF baseline to medication-related daytime beta modulation under continuous closed-loop stimulation adjustment.

During online operation, the implantable pulse generator recorded the smoothed beta-energy biomarker used by the controller and the stimulation amplitude delivered at each control step. Device timestamps were synchronized with external clock time, allowing the recorded biomarker and stimulation sequences to be aligned with medication intake and other experimental events. After each session, the data stored in the implantable pulse generator were retrieved through Bluetooth communication.

These embedded recordings were used to evaluate whether the controller executed the intended stimulation-update logic under real patient-in-the-loop conditions. Specifically, we assessed whether the recorded stimulation amplitudes remained within the predefined safety range, whether single-step amplitude changes satisfied the maximum update constraint, and whether the stored stimulation sequence was consistent with the controller decision rule when replayed offline using the recorded beta-energy sequence. This analysis was designed to evaluate embedded execution consistency and constraint satisfaction, not clinical efficacy.

S3. Beta preprocessing and Patient-Specific Calibration

S3.1 Beta-energy feature extraction

For offline model calibration, STN LFP recordings were processed in 1-min windows to match the minute-scale control update interval. For each window, the power spectral density was estimated using short-time Fourier transform with a Hanning window. Beta energy was defined as the mean power spectral density within the 12–30 Hz band. To reduce the influence of global power fluctuations and inter-recording amplitude differences, beta energy was normalized by the mean power spectral density within the 5–55 Hz band:

$$B_{k}^{raw}=\frac{\mathrm{mean}\left[ PSD_{k}(f), f\in12\text{–}30\mathrm{Hz} \right]}{\mathrm{mean}\left[ PSD_{k}(f), f\in5\text{–}55\mathrm{Hz} \right]}.$$

Here, $k$denotes the 1-min analysis window. This normalized beta-energy feature was used for offline stimulation–beta calibration, medication-cycle analysis, fast–slow dynamic fitting, and noise estimation.

During online embedded closed-loop operation, beta energy was computed by the implantable pulse generator using FFT-based spectral estimation with a Hanning window. The same beta-band definition was used for online control, with beta energy derived from the 12–30 Hz band and normalized by the broadband 5–55 Hz power. The implantable pulse generator stored the smoothed beta-energy feature used by the controller, rather than continuous raw LFP. Therefore, offline calibration analyses were based on post-processed LFP-derived beta energy at 1-min resolution, whereas online replay analyses used the device-stored smoothed beta energy and stimulation amplitude recorded at each control step.

In addition, the embedded FFT-based beta-energy calculation was cross-validated against the offline MATLAB implementation using the same input LFP segments and spectral settings. The beta-energy values computed by the implantable pulse generator and by MATLAB agreed within machine-precision error, confirming numerical consistency between the embedded and offline feature-extraction pipelines.

S3.2 Patient-specific steady-state stimulation–beta map calibration

For each patient, the steady-state stimulation–beta relationship was calibrated from the MED OFF stimulation-amplitude modulation recordings. At each stimulation amplitude, the 5-min steady-state segment following the 10-min wash-in period was used to estimate the amplitude-specific beta equilibrium. The beta-energy values within this steady-state segment were averaged to obtain $B_{\mathrm{recorded}}^{\left( i \right)}(I_{m})$, where $I_{m}$denotes the $m$-th stimulation amplitude and $i$denotes the patient index.

The computational basal ganglia prior described in Section S1 was used only as a shape constraint for the stimulation–beta relationship. This prior was aligned to the recorded steady-state beta values of each patient using patient-specific scaling and offset parameters, yielding the calibrated steady-state map $B_{\infty}^{\left( i \right)}(I)$. The resulting map was used as the equilibrium term in the minute-scale beta model and as the source of patient-specific stimulation sensitivity for the embedded controller.

The fitting error was quantified as:

$$\mathrm{RMSE}_{\mathrm{ss}}^{\left( i \right)}=\sqrt{\frac{1}{N_{I}}\sum_{m=1}^{N_{I}} \left[ B_{\mathrm{recorded}}^{\left( i \right)}(I_{m})-B_{\infty}^{\left( i \right)}(I_{m}) \right]^{2}},$$

where $N_{I}$is the number of stimulation amplitudes tested in patient $i$.

S3.3 Fast–slow dynamics after stimulation changes

The patient-specific steady-state map $B_{\infty}^{\left( i \right)}(I)$defines the beta-energy equilibrium expected under a fixed stimulation amplitude. However, after a stimulation-amplitude update, beta activity did not instantaneously reach the new equilibrium. Therefore, stimulation-induced beta dynamics were modeled using a fast–slow approximation that separates the immediate response to the new stimulation amplitude from the subsequent minute-scale relaxation process.

For each patient, transient beta responses were estimated from the 10-min wash-in segment following each stimulation-amplitude change during the MED OFF amplitude-modulation recordings. The 5-min steady-state segment after the wash-in period was used for steady-state map calibration as described in Section S3.2, whereas the wash-in segment was used to estimate the transient response parameters.

The slow stimulation-related beta state was modeled as a first-order relaxation process toward the patient-specific steady-state beta value:

$$\tau_{s}^{\left( i \right)}\frac{dB_{s}^{\left( i \right)}(t)}{dt}=B_{\infty}^{\left( i \right)}(I(t))-B_{s}^{\left( i \right)}(t),$$

where $B_{s}^{\left( i \right)}(t)$denotes the slow stimulation-related beta state, $B_{\infty}^{\left( i \right)}(I(t))$is the patient-specific steady-state beta value predicted by the calibrated stimulation–beta map, and $\tau_{s}^{\left( i \right)}$is the patient-specific relaxation time constant.

To capture the immediate beta change after stimulation adjustment, the stimulation-related beta component was represented as a weighted combination of the current steady-state prediction and the slow relaxation state:

$$B_{\mathrm{stim},k}^{\left( i \right)}=w_{i}B_{\infty}^{\left( i \right)}(I_{k})+(1-w_{i})B_{s,k}^{\left( i \right)},$$

where $w_{i}$denotes the fast jump weight. This formulation allows the beta trajectory to exhibit an immediate shift after stimulation amplitude changes, followed by gradual convergence toward the new equilibrium over subsequent control steps.

The transient parameters $k_{i}$and $w_{i}$were estimated by minimizing the discrepancy between the recorded wash-in beta trajectory and the simulated beta response after each stimulation-amplitude change. The fitting error was quantified as:

$$\mathrm{RMSE}_{\mathrm{dyn}}^{\left( i \right)}=\sqrt{\frac{1}{N_{T}}\sum_{t=1}^{N_{T}} \left[ B_{\mathrm{recorded}}^{\left( i \right)}(t)-B_{\mathrm{simulated}}^{\left( i \right)}(t) \right]^{2}},$$

where $N_{T}$is the number of 1-min samples included in the wash-in segments. In cases where direct fitting of $k_{i}$was poorly identified or reached the fitting boundary, the relaxation parameter was replaced by the value estimated from transition/autocorrelation dynamics. These cases are indicated in Supplementary Table S9.

Together, $B_{\infty}^{\left( i \right)}(I)$, $k_{i}$, and $w_{i}$defined the patient-specific stimulation-response dynamics used in the closed-loop simulations. The calibrated fast–slow parameters, fitting errors, and fitting notes are summarized together with the other patient-specific calibration parameters in Supplementary Table S9.

S3.4 Patient-specific target-zone definition

For each patient, the target beta zone was derived from the calibrated steady-state stimulation–beta map $B_{\infty}^{\left( i \right)}(I)$. The clinically tolerated stimulation working range was first determined and denoted as $\left[ I_{L}^{\left( i \right)} , I_{H}^{\left( i \right)} \right]$. Here, $I_{L}^{\left( i \right)}$and $I_{H}^{\left( i \right)}$represent the lower and upper stimulation amplitudes allowed during closed-loop control.

The calibrated steady-state map was evaluated at both stimulation boundaries. Because beta energy generally decreases with increasing stimulation amplitude, $B_{\infty}^{\left( i \right)}(I_{H}^{\left( i \right)})$represents the lower beta steady-state value, whereas $B_{\infty}^{\left( i \right)}(I_{L}^{\left( i \right)})$represents the higher beta steady-state value. To avoid defining the control zone exactly at the clinical stimulation extremes, the target-zone boundaries were placed inside the beta range spanned by these two steady-state values:

$$B_{L}^{\left( i \right)}=0.95B_{\infty}^{\left( i \right)}(I_{H}^{\left( i \right)})+0.05B_{\infty}^{\left( i \right)}(I_{L}^{\left( i \right)}),$$

$$B_{H}^{\left( i \right)}=0.5B_{\infty}^{\left( i \right)}(I_{H}^{\left( i \right)})+0.5B_{\infty}^{\left( i \right)}(I_{L}^{\left( i \right)}).$$

Thus, $\left[ B_{L}^{\left( i \right)} , B_{H}^{\left( i \right)} \right]$was defined individually for each patient from the calibrated stimulation–beta relationship and the clinically tolerated stimulation working range, rather than from a fixed group-level beta threshold. Medication-related beta modulation was modeled separately and was not used to define the target-zone boundaries.

S3.5 Estimation of endogenous fluctuation, observation noise, and background drift

The minute-scale $\beta_{\mathrm{STN}}$model separated non-stimulation-related variability into three components: endogenous fluctuation of the latent beta state, observation noise in the biomarker available to the controller, and slower background drift. This separation was used to distinguish intrinsic beta variability from measurement-level disturbances that directly affect feedback decisions.

For each patient, residual beta fluctuations were estimated after removing the stimulation-related component and levodopa-related modulation from the recorded beta-energy sequence. Specifically, the calibrated stimulation-related term $B_{\mathrm{stim}}^{\left( p \right)}(k)$was first computed from the patient-specific stimulation–$\beta_{\mathrm{STN}}$ map and fast–slow stimulation-response model. The levodopa-related modulation term $B_{\mathrm{LD}}^{\left( p \right)}(k)$was then estimated from the shared medication-response temporal profile and the patient-specific modulation amplitude. The residual sequence was calculated as

$$r^{\left( p \right)}(k)=B_{\mathrm{rec}}^{\left( p \right)}(k)-B_{\mathrm{stim}}^{\left( p \right)}(k)-B_{\mathrm{LD}}^{\left( p \right)}(k),$$

where $B_{\mathrm{rec}}^{\left( p \right)}(k)$denotes the recorded normalized beta-energy sequence of patient $p$at the $k$-th minute-scale sample. This residual sequence contains unresolved endogenous beta variability, slow background modulation, and observation-level measurement disturbance.

To estimate endogenous fluctuation, the residual sequence was decomposed into a slow component and a remaining high-frequency component. The slow residual variation was treated as process-level fluctuation acting on the latent beta state. Its innovation term was modeled as zero-mean Gaussian noise:

$$\xi_{s}^{\left( p \right)}(k)\sim\mathcal{N}\left( 0 , \sigma_{\xi,p}^{2} \right),$$

where $\sigma_{\xi,p}$denotes the patient-specific endogenous fluctuation strength. This term was added to the slow stimulation-related beta state in the fast–slow dynamical model:

$$\tau_{s,p}\frac{dB_{s}^{\left( p \right)}(t)}{dt}=B_{\infty}^{\left( p \right)}(I(t))-B_{s}^{\left( p \right)}(t)+\xi_{s}^{\left( p \right)}(t).$$

Thus, endogenous fluctuation affected the latent beta trajectory $B_{\mathrm{true}}^{\left( p \right)}(t)$and was included in formal performance evaluation.

Observation noise was estimated from the high-frequency component of the residual sequence after removal of the slow component. This term represents measurement-level variability caused by spectral-estimation error, stimulation-related recording contamination, movement artifacts, electrocardiographic artifacts, and other disturbances in the controller-visible biomarker. Observation noise was modeled as

$$\epsilon_{\mathrm{obs}}^{\left( p \right)}(k)\sim\mathcal{N}\left( 0 , \sigma_{\epsilon,p}^{2} \right),$$

where $\sigma_{\epsilon,p}$denotes the patient-specific observation-noise standard deviation. Unlike $\xi_{s}^{\left( p \right)}(k)$, $\epsilon_{\mathrm{obs}}^{\left( p \right)}(k)$was added only to the observed biomarker:

$$B_{\mathrm{obs}}^{\left( p \right)}(k)=B_{\mathrm{true}}^{\left( p \right)}(k)+\epsilon_{\mathrm{obs}}^{\left( p \right)}(k).$$

Therefore, observation noise affected controller decisions but did not alter the latent beta state used for formal performance evaluation.

In addition to Gaussian observation noise, a sinusoidal observation disturbance was introduced as a stress-test condition to evaluate controller behavior under structured non-stationary sensing disturbance. This term was not intended to represent a fitted physiological rhythm. Instead, it was used to simulate a strong periodic artifact or slowly varying observation bias. The sinusoidal disturbance was defined as

$$\epsilon_{\sin}^{\left( p \right)}(k)=A_{\sin,p}\sin\left( \omega k\Delta t + \phi\right),$$

where

$$A_{\sin,p}=2.5\sigma_{\epsilon,p},$$

and

$$\omega=\frac{2\pi}{T_{\sin}}.$$

Here, $\Delta t=1\text{ }\min$, $T_{\sin}=480\text{ }\min$, and $\phi$denotes the initial phase, which was randomized across repeated simulations. In the sinusoidal-disturbance scenario, the controller-visible biomarker was defined as

$$B_{\mathrm{obs}}^{\left( p \right)}(k)=B_{\mathrm{true}}^{\left( p \right)}(k)+\epsilon_{\sin}^{\left( p \right)}(k).$$

Slow background drift, denoted as $B_{\mathrm{bg}}^{\left( p \right)}(t)$, was included in the general model definition to represent unmodeled long-timescale changes in beta activity that may arise from circadian state, arousal, behavioral context, or gradual baseline shifts. In the formal four-scenario controller comparisons, $B_{\mathrm{bg}}^{\left( p \right)}(t)$was not introduced as an additional independent disturbance term because the simulation window focused on an 8-h daytime medication-related control period, whereas background drift mainly reflects slower circadian-scale modulation. Its empirical contribution was therefore absorbed into the estimated medication/background modulation and endogenous fluctuation terms. Accordingly, the formal disturbance scenarios separately tested levodopa-related modulation, endogenous process fluctuation, Gaussian observation noise, and sinusoidal observation disturbance, without adding an independent circadian-drift module.

The estimated patient-specific values of $\sigma_{\xi,p}$and $\sigma_{\epsilon,p}$are summarized in Supplementary Table S9. These values were fixed for each patient and used consistently across the closed-loop simulations, ablation analyses, and parameter-mismatch tests.

TABLE S9. Patient-specific calibration parameters used for simulation and embedded control

| **Parameter** | **P1** | **P2** | **P3** | **P4** | **P5** |
| --- | --- | --- | --- | --- | --- |
| $I_{L}$(mA) | 2.05 | 1.80 | 2.95 | 1.25 | 1.25 |
| $I_{H}$(mA) | 2.55 | 2.10 | 3.25 | 1.40 | 1.45 |
| Calibration range (mA) | 1.94–2.81 | 1.614–2.136 | 2.922–3.328 | 1.234–1.466 | 1.23–1.52 |
| $I_{0}$(mA) | 2.35 | 1.85 | 2.95 | 1.25 | 1.40 |
| $a_{p}$ | 0.2423 | 0.4009 | 0.3183 | 1.8539 | 2.5120 |
| $b_{p}$ | 0.8848 | 1.9392 | 1.4584 | 1.0628 | 0.9633 |
| $s_{p}$ | 0.2226 | 0.3555 | 0.6638 | 7.4312 | 7.7390 |
| $B_{\infty}(I_{L})$ | 1.0176 | 2.0409 | 1.7048 | 2.4978 | 2.9077 |
| $B_{\infty}(I_{H})$ | 0.8914 | 1.9397 | 1.4603 | 1.1065 | 0.9975 |
| $B_{L}$ | 0.8977 | 1.9447 | 1.4725 | 1.1761 | 1.0930 |
| $B_{H}$ | 0.9545 | 1.9903 | 1.5825 | 1.8022 | 1.9526 |
| $A_{\mathrm{med}}$ | 0.2546 | 0.2495 | 0.3051 | 1.6238 | 1.9455 |
| $\tau_{s}$(min) | 18.2434 | 14.2526 | 22.9745 | 18.5714 | 19.6494 |
| $w_{p}$ | 0.0465 | 0.0321 | 0.0177 | 0.0088 | 0.0184 |
| $\sigma_{\mathrm{obs}}$ | 0.0462 | 0.0975 | 0. 0858 | 0. 0072 | 0. 0051 |
| $\sigma_{\mathrm{endo}}$ | 0.0171 | 0.0545 | 0.0119 | 0.0367 | 0.0389 |
| Fit RMSE | 0.1790 | 0.3211 | 0.1865 | 0.3082 | 1.7209 |

**Table note.** $I_{L}$and $I_{H}$denote the lower and upper stimulation amplitudes allowed during online control. The calibration range denotes the full stimulation range used for patient-specific stimulation–beta map fitting. $I_{0}$is the initial stimulation amplitude. $a_{p}$, $b_{p}$, and $s_{p}$are patient-specific alignment parameters used to calibrate the computational steady-state stimulation–beta prior to the recorded stimulation–beta data. $B_{\infty}(I_{L})$and $B_{\infty}(I_{H})$are the steady-state beta values predicted by the calibrated stimulation–beta map at the lower and upper stimulation boundaries, respectively. Because beta energy generally decreases with increasing stimulation amplitude, $B_{\infty}(I_{H})$is lower than $B_{\infty}(I_{L})$. $B_{L}$and $B_{H}$denote the lower and upper boundaries of the patient-specific beta target zone. $A_{\mathrm{med}}$denotes the medication/background modulation amplitude. $\tau_{s}$is the minute-scale relaxation time constant after stimulation-amplitude changes, and $w_{p}$ is the fast jump weight. $\sigma_{\mathrm{obs}}$denotes the observation-noise standard deviation, whereas $\sigma_{\mathrm{endo}}$denotes the endogenous beta fluctuation strength. Fit RMSE quantifies the fitting error of the patient-specific stimulation–beta calibration. All beta-related quantities are normalized beta-energy values.

S4. Responses of *β*_STN_ to Levodopa intake

Responses of $\beta_{\mathrm{STN}}$to levodopa intake were modeled from medication-cycle recordings. Participants withheld dopaminergic medication for at least 12 h before the experiment. Continuous STN LFP was recorded from 30 min before medication intake to 120 min after medication intake, while stimulation parameters were kept unchanged. The 30-min pre-medication segment was used as the MED OFF baseline.

The rate of change in systemic levodopa concentration was modeled using a one-compartment absorption–elimination process. The absorbed drug amount $A$is influenced by the absorption rate constant $k_{a}$and the elimination rate constant $\lambda$:

$$\frac{dA}{dt}=k_{a}A_{a}-\lambda A,$$

where $A_{a}$is the amount of drug remaining to be absorbed and $A$is the absorbed drug amount.

Solving this differential equation gives:

$$A=A_{0}\frac{k_{a}}{k_{a}-\lambda}\left( e^{-\lambda t} - e^{-k_{a}t} \right),$$

where $A_{0}$denotes the initial levodopa dose. Therefore, the systemic levodopa concentration can be expressed as:

$$c_{\mathrm{med}}=\frac{F}{V}A=\frac{FA_{0}}{V}\frac{k_{a}}{k_{a}-\lambda}\left( e^{-\lambda t} - e^{-k_{a}t} \right)=C_{0}^{'}\left( e^{-\lambda t} - e^{-k_{a}t} \right),$$

where

$$C_{0}^{'}=\frac{FA_{0}}{V}\frac{k_{a}}{k_{a}-\lambda}.$$

Here, $F$is bioavailability, $V$is distribution volume, $k_{a}$is the absorption rate constant, and $\lambda$is the elimination rate constant. Because plasma levodopa concentration was not directly measured in this study, $C_{0}^{'}$was absorbed into the subsequent scaling parameters.

The dynamic response of $\beta_{\mathrm{STN}}$to levodopa intake was modeled as a sigmoid transformation of the absorption–elimination time course:

$$\beta_{\mathrm{STN},\mathrm{med}}=f(c_{\mathrm{med}})=M_{1}\frac{1}{1+e^{-M_{3}(c_{\mathrm{med}}+M_{2})}}+M_{4}.$$

Substituting the concentration model into this equation gives:

$$\beta_{\mathrm{STN},\mathrm{med}}=M_{1}\frac{1}{1+e^{-M_{3}\left[ C_{0}^{'}\left( e^{-\lambda t} - e^{-k_{a}t} \right) + M_{2} \right]}}+M_{4}.$$

Since $C_{0}^{'}$was not separately identifiable without direct plasma concentration measurements, the model was reparameterized as:

$$\beta_{\mathrm{STN},\mathrm{med}}=M_{1}\frac{1}{1+e^{-M_{3}^{'}\left[ e^{-\lambda t}-e^{-k_{a}t}+M_{2}^{'} \right]}}+M_{4}.$$

Here, $M_{2}^{'}$, $M_{3}^{'}$, $M_{4}$, $k_{a}$, and $\lambda$define the shared temporal shape of levodopa-related beta modulation, whereas $M_{1}$determines the modulation amplitude.

In the revised multi-patient calibration, the medication-response shape parameters $M_{2}^{'}$, $M_{3}^{'}$, $M_{4}$, $k_{a}$, and $\lambda$were estimated by pooled fitting across medication-cycle recordings from all five patients. This shared-shape fitting strategy was used to reduce overfitting of pharmacokinetic shape parameters at the individual-patient level. Patient-specific variability in levodopa-related beta modulation was represented by the amplitude term $M_{1}^{\left( i \right)}$, denoted as $A_{\mathrm{med}}^{\left( i \right)}$in the simulation model:

$$\beta_{\mathrm{STN},\mathrm{med}}^{\left( i \right)}(t)=A_{\mathrm{med}}^{\left( i \right)}\frac{1}{1+e^{-M_{3}^{'}\left[ e^{-\lambda t}-e^{-k_{a}t}+M_{2}^{'} \right]}}+M_{4}.$$

Equivalently, the shared medication temporal profile can be written as:

$$\phi_{\mathrm{med}}(t)=\frac{1}{1+e^{-M_{3}^{'}\left[ e^{-\lambda t}-e^{-k_{a}t}+M_{2}^{'} \right]}},$$

and the patient-specific levodopa-related beta modulation is:

$$\beta_{\mathrm{STN},\mathrm{med}}^{\left( i \right)}(t)=A_{\mathrm{med}}^{\left( i \right)}\phi_{\mathrm{med}}(t)+M_{4}.$$

Bayesian optimization was used to estimate the shared medication-response parameters, using MATLAB’s built-in bayesopt function (R2022b, The MathWorks, Natick, MA). The objective function minimized the RMSE between simulated and recorded medication-cycle beta trajectories across all included patients. The optimization process was configured with a maximum of 350 objective evaluations and used the acquisition function expected-improvement-plus to balance exploration and exploitation during parameter search. The shared medication-response parameters are summarized in Supplementary Table S10. Patient-specific modulation amplitudes $A_{\mathrm{med}}^{\left( i \right)}$are listed in Supplementary Table S9.

TABLE S10 Shared parameters of levodopa-related beta modulation

| $\boldsymbol{M}_{\boldsymbol{2}}^{\boldsymbol{'}}$ | $\boldsymbol{M}_{\boldsymbol{3}}^{\boldsymbol{'}}$ | $\boldsymbol{M}_{\boldsymbol{4}}$ | $\boldsymbol{\lambda}$ | $\boldsymbol{k}_{\boldsymbol{a}}$ |
| --- | --- | --- | --- | --- |
| 0.0502 | -298.6 | 0.8 | 0.01367 | 0.01139 |

S5. Derivation of the embedded explicit TZPC control law

The full trend–zone predictive control problem evaluates the predicted beta value after candidate stimulation updates and selects the update that minimizes a zone-based objective. To reduce online computational burden, this problem was locally linearized around the current stimulation working point and transformed into an explicit feedback law.

Let

$$d_{k}=\hat{\Delta B}_{k}$$

denote the estimated local beta trend at control step $k$. The no-action one-step prediction, i.e., the predicted beta value without applying a new stimulation update, was defined as

$$\hat{B}_{k+1\mid k}^{0}=\tilde{B}_{k}+\lambda_{\Delta}d_{k},$$

where $\tilde{B}_{k}$is the smoothed beta-energy value and $\lambda_{\Delta}$is the trend extrapolation weight.

Around the current stimulation working point, the patient-specific action effect was locally linearized as

$$G_{i}(u_{k};I_{k})\approx-\eta_{i}u_{k},\eta_{i}>0,$$

where $u_{k}$is the stimulation-amplitude update and $\eta_{i}$is the local stimulation sensitivity of patient $i$. Thus, the predicted beta value after applying $u_{k}$was

$$\hat{B}_{k+1\mid k}(u_{k})=\hat{B}_{k+1\mid k}^{0}-\eta_{i}u_{k},$$

or equivalently,

$$\hat{B}_{k+1\mid k}(u_{k})=\tilde{B}_{k}+\lambda_{\Delta}d_{k}-\eta_{i}u_{k}.$$

The theoretical one-step TZPC objective was defined as

$$J_{k}(u_{k})=q_{z}D^{2}\left( \hat{B}_{k+1\mid k}(u_{k}),[B_{L},B_{H}] \right)+q_{v}\left( \hat{B}_{k+1\mid k}(u_{k})-\tilde{B}_{k} \right)^{2}+ru_{k}^{2},$$

where $q_{z}$, $q_{v}$, and $r$are the zone-violation penalty, predicted beta-change penalty, and stimulation-update penalty, respectively. The squared distance to the target zone was defined as

$$D^{2}(x,[B_{L},B_{H}])=[\max(0,x-B_{H})]^{2}+[\max(0,B_{L}-x)]^{2}.$$

The no-action zone error was defined as

$$E_{k}=\hat{B}_{k+1\mid k}^{0}-\mathrm{clip}\left( \hat{B}_{k+1\mid k}^{0} , B_{L} , B_{H} \right).$$

Therefore,

$$E_{k}=0,\hat{B}_{k+1\mid k}^{0}\in[B_{L},B_{H}],$$

$$E_{k}>0,\hat{B}_{k+1\mid k}^{0}>B_{H},$$

$$E_{k}<0,\hat{B}_{k+1\mid k}^{0}<B_{L}.$$

We further defined an activation variable for the zone-error term:

$$\sigma_{k}=\left\{ \begin{matrix} 0, & E_{k}=0, \\ 1, & E_{k}\neq0. \end{matrix} \right.$$

Under a local active-set approximation, the zone-distance term can be approximated as

$$D^{2}\left( \hat{B}_{k+1\mid k}(u_{k}),[B_{L},B_{H}] \right)\approx\sigma_{k}(E_{k}-\eta_{i}u_{k})^{2}.$$

Meanwhile,

$$\hat{B}_{k+1\mid k}(u_{k})-\tilde{B}_{k}=\lambda_{\Delta}d_{k}-\eta_{i}u_{k}.$$

Thus, the local quadratic objective becomes

$$J_{k}^{\mathrm{loc}}(u_{k})=q_{z}\sigma_{k}(E_{k}-\eta_{i}u_{k})^{2}+q_{v}(\lambda_{\Delta}d_{k}-\eta_{i}u_{k})^{2}+ru_{k}^{2}.$$

Taking the derivative with respect to $u_{k}$gives

$$\frac{\partial J_{k}^{\mathrm{loc}}}{\partial u_{k}}=-2q_{z}\sigma_{k}\eta_{i}(E_{k}-\eta_{i}u_{k})-2q_{v}\eta_{i}(\lambda_{\Delta}d_{k}-\eta_{i}u_{k})+2ru_{k}.$$

Setting the derivative to zero yields

$$-2q_{z}\sigma_{k}\eta_{i}E_{k}+2q_{z}\sigma_{k}\eta_{i}^{2}u_{k}-2q_{v}\eta_{i}\lambda_{\Delta}d_{k}+2q_{v}\eta_{i}^{2}u_{k}+2ru_{k}=0.$$

Rearranging terms gives

$$\left[ (q_{v}+q_{z}\sigma_{k})\eta_{i}^{2}+r \right]u_{k}=q_{v}\eta_{i}\lambda_{\Delta}d_{k}+q_{z}\sigma_{k}\eta_{i}E_{k}.$$

Therefore, the unconstrained explicit stimulation command is

$$u_{k}^{\mathrm{cmd}}=\frac{q_{v}\eta_{i}\lambda_{\Delta}}{(q_{v}+q_{z}\sigma_{k})\eta_{i}^{2}+r}d_{k}+\frac{q_{z}\sigma_{k}\eta_{i}}{(q_{v}+q_{z}\sigma_{k})\eta_{i}^{2}+r}E_{k}.$$

Since $d_{k}=\hat{\Delta B}_{k}$, this can be written as

$$u_{k}^{\mathrm{cmd}}=K_{\Delta,i}^{\left( \sigma_{k} \right)}\hat{\Delta B}_{k}+K_{Z,i}^{\left( \sigma_{k} \right)}E_{k},$$

where

$$K_{\Delta,i}^{\left( \sigma_{k} \right)}=\frac{q_{v}\eta_{i}\lambda_{\Delta}}{(q_{v}+q_{z}\sigma_{k})\eta_{i}^{2}+r},$$

and

$$K_{Z,i}^{\left( \sigma_{k} \right)}=\frac{q_{z}\sigma_{k}\eta_{i}}{(q_{v}+q_{z}\sigma_{k})\eta_{i}^{2}+r}.$$

When the no-action prediction remains inside the target zone,

$$E_{k}=0,\sigma_{k}=0,$$

and the update reduces to a pure trend-driven command:

$$u_{k}^{\mathrm{cmd}}=K_{\Delta,i}^{\mathrm{in}}\hat{\Delta B}_{k},$$

with

$$K_{\Delta,i}^{\mathrm{in}}=\frac{q_{v}\eta_{i}\lambda_{\Delta}}{q_{v}\eta_{i}^{2}+r},$$

and

$$K_{Z,i}^{\mathrm{in}}=0.$$

When the no-action prediction is outside the target zone,

$$E_{k}\neq0,\sigma_{k}=1,$$

the update becomes

$$u_{k}^{\mathrm{cmd}}=K_{\Delta,i}^{\mathrm{out}}\hat{\Delta B}_{k}+K_{Z,i}^{\mathrm{out}}E_{k},$$

where

$$K_{\Delta,i}^{\mathrm{out}}=\frac{q_{v}\eta_{i}\lambda_{\Delta}}{(q_{v}+q_{z})\eta_{i}^{2}+r},$$

and

$$K_{Z,i}^{\mathrm{out}}=\frac{q_{z}\eta_{i}}{(q_{v}+q_{z})\eta_{i}^{2}+r}.$$

Finally, the stimulation update was constrained by the maximum single-step change and the stimulation safety range. The constrained update was

$$u_{k}^{*}=\mathrm{clip}\left( u_{k}^{\mathrm{cmd}},-\Delta I_{\max},+\Delta I_{\max} \right),$$

and the next stimulation amplitude was

$$I_{k+1}=\mathrm{clip}\left( I_{k}+u_{k}^{*},I_{\min},I_{\max} \right).$$

For embedded implementation, the update was further quantized to the stimulation amplitude resolution $\delta_{I}$:

$$u_{k}^{*,q}=\delta_{I}\cdot\mathrm{round}\left( \frac{u_{k}^{*}}{\delta_{I}} \right),$$

$$I_{k+1}=\mathrm{clip}\left( I_{k}+u_{k}^{*,q},I_{\min},I_{\max} \right).$$

Thus, the embedded TZPC controller can be implemented as a fixed explicit trend–zone feedback law:

$$u_{k}^{\mathrm{cmd}}=K_{\Delta,i}\hat{\Delta B}_{k}+K_{Z,i}E_{k},$$

followed by step-size clipping, amplitude quantization, and stimulation-range clipping. Online execution therefore requires only smoothing, trend estimation, zone-error calculation, scalar multiplication, and bound checking, without iterative optimization or online nonlinear model evaluation.

In addition, because the stimulation update must ultimately be quantized by $Q_{0.025}(\cdot)$, an excessively small zone-correction command may be quantized to 0. To avoid forming an inactive dead zone near the original target-zone boundaries, eTZPC used a slightly inward-contracted control zone during online decision making:

$$[B_{L,i}^{c},B_{H,i}^{c}]=\left[ B_{L}+\frac{\Delta I_{\mathrm{res}}}{2K_{Z,i}},\text{ }B_{H}-\frac{\Delta I_{\mathrm{res}}}{2K_{Z,i}} \right].$$

This contracted zone was used only for online control decisions, whereas formal performance evaluation was still based on the original target zone $\left[ B_{L} , B_{H} \right]$.

S6. Baseline controllers and ablation variants

All controller-parameter optimization and closed-loop simulations were implemented in MATLAB R2022b (The MathWorks, Natick, MA, USA). The final fixed controller parameters used in the formal evaluation are summarized in Supplementary Table S11.

For each candidate parameter set, the controller was run in the parameter-optimization environment, and a composite loss function was calculated:

$$L=0.8\cdot\mathrm{TIR}+2.5\cdot(1-\mathrm{TIR})+0.015\cdot\mathrm{SwitchingBurden}+0.05\cdot AverageI.$$

Here, $\mathrm{TimeInRange}$ denotes the proportion of time during which the latent $\beta_{\mathrm{STN}}$remained within the target zone, $\mathrm{ViolationArea}$denotes the cumulative zone-violation area after the latent $\beta_{\mathrm{STN}}$exceeded the target zone, $\mathrm{SwitchingBurden}$denotes the number of stimulation updates or cumulative switching burden, and $\mathrm{AverageI}$denotes the mean stimulation amplitude. The complete definitions of these metrics are provided in the Control Performance Metrics subsection of the main text.

The final controller parameters selected from the tuning environment and kept fixed in all formal evaluations are summarized in Supplementary Table S11.

TABLE S11. Fixed controller parameters

| **eTZPC parameter** | $q_{z}$ | $q_{v}$ | $r$ | $\lambda_{\Delta}$ | $\rho$ | $\Delta I_{\max}$ | $\delta_{I}$ |
| --- | --- | --- | --- | --- | --- | --- | --- |
| Value | 2.2000 | 3.5422 | 0.001085 | 1.0234 | 0.125 | 0.10 mA | 0.025 mA |
| **Baseline parameter** | $\Delta I_{\mathrm{DT}}$ | $K_{P}$ | $K_{I}$ | $K_{D}$ |  |  |  |
| Value | 0.05 mA | 2.5277 | $9.1814\times{10}^{-7}$ | 0.035679 |  |  |  |

**Table note.** Parameters were fixed for all formal evaluations. $q_{z}$, $q_{v}$, $r$, and $\lambda_{\Delta}$are eTZPC parameters; $\rho$is the smoothing coefficient; $\Theta_{\Delta}$is the trend deadband; $\Delta I_{\max}$and $\delta_{I}$define stimulation update constraints. $\Delta I_{\mathrm{DT}}$is the dual-threshold update step, and $K_{P}$, $K_{I}$, and $K_{D}$are PID gains.

S7 Progressive Ablation Analysis

To clarify the contribution of each component of eTZPC, a progressive ablation analysis was performed. Starting from the most basic reactive dual-threshold controller, observation smoothing, trend prediction, patient-specific action prediction, and embedded explicit implementation were sequentially added. Each step corresponded to a specific closed-loop failure mode and was used to test the effect of that module on false triggering, feedback delay, individualized mismatch, and online complexity.

All ablation variants used the same patient model, target zone, stimulation safety boundaries, maximum step-size constraint, and performance metrics. The variants were defined as follows.

**A0: Current-value dual-threshold control.**
This variant directly adjusted stimulation according to whether the current observation $B_{\mathrm{obs},k}$exceeded the target zone. It did not include observation smoothing, trend prediction, or model-based action effects, and represented the most basic reactive threshold controller.

**A1: A0 + observation smoothing.**
This variant first recursively smoothed $B_{\mathrm{obs},k}$, and then performed dual-threshold judgment based on $\tilde{B}_{k}$. This variant was used to evaluate whether smoothing reduced false triggering and frequent stimulation switching caused by random observation noise.

**A2: A1 + trend-based no-action prediction.**
This variant further calculated the local trend,

$$\hat{\Delta B}_{k}=\tilde{B}_{k}-\tilde{B}_{k-1},$$

and constructed a one-step no-action prediction:

$$\hat{B}_{k+1\mid k}^{0}=\tilde{B}_{k}+\lambda\hat{\Delta B}_{k}.$$

The controller performed threshold judgment based on $\hat{B}_{k+1\mid k}^{0}$, rather than the current $\tilde{B}_{k}$. This variant was used to evaluate whether trend prediction reduced the feedback delay of reactive threshold control.

**A3: A2 + patient-specific nonlinear action prediction.**
This variant used the patient-specific action effect $G_{i}(u_{k};I_{k})$to calculate the predicted value after candidate stimulation updates, and minimized the TZPC zone-predictive cost within a finite action set:

$$u_{k}^{*}=\arg\min_{u_{k}\in\mathcal{U}_{k}}J_{k}(u_{k}).$$

This variant corresponded to the full nonlinear TZPC and was used to evaluate the contribution of the patient-specific stimulation–response model and zone-predictive cost to control performance.

**A4: Embedded explicit implementation of A3.**
This variant compressed the nonlinear action effect $G_{i}(u_{k};I_{k})$into the linear approximation $G_{i}(u_{k};I_{k})\approx-\eta_{i}u_{k}$, and adopted the explicit trend–zone feedback law:

$$u_{k}^{\mathrm{cmd}}=K_{\Delta,i}\hat{\Delta B}_{k}+K_{Z,i}E_{k}.$$

This variant was the final online eTZPC and was used to evaluate whether local linearization and fixed-gain implementation preserved control performance close to the full TZPC while reducing online computational complexity.

Through this progressive design, A0–A4 respectively tested the contributions of reactive threshold control, observation denoising, anticipatory trend response, patient-calibrated action prediction, and embedded reduced-order implementation to closed-loop performance.

The ablation variants are summarized in Supplementary Table S12.

TABLE S12. Ablation variants

| **Variant** | **Definition** | **Main component tested** |
| --- | --- | --- |
| A0 | Current-value DT | Reactive threshold baseline |
| A1 | A0 + smoothing | Noise-driven false triggering |
| A2 | A1 + trend prediction | Feedback delay under directional beta changes |
| A3 | A2 + patient-specific action prediction | Patient-specific stimulation–beta response |
| A4 | Embedded eTZPC | Explicit embedded implementation and switching stabilization |

S8 Statistical Analysis

All performance metrics were first calculated within each individual closed-loop simulation. For repeated simulations under the same patient, test scenario, and controller, results obtained from different random noise realizations and initial disturbance phases were first averaged to generate patient-level performance values. Results were then summarized at the patient level and reported as mean ± standard deviation. This procedure avoided treating repeated simulation runs as independent clinical samples.

Comparisons among different controllers used a paired design. The performance of eTZPC, dual-threshold control, and PID control was compared within the same patient, test scenario, and disturbance realization. For each test scenario and each primary metric, the Friedman test was first used to evaluate the overall difference among the three controllers. If the overall test was significant, pairwise comparisons were then performed using the paired Wilcoxon signed-rank test, with Holm–Bonferroni correction for multiple comparisons.

The progressive ablation analysis used predefined adjacent-step comparisons, namely A0 vs. A1, A1 vs. A2, A2 vs. A3, and A3 vs. A4, to evaluate the incremental contributions of observation smoothing, trend prediction, patient-specific action prediction, and embedded explicit implementation, respectively. Adjacent steps were compared using the paired Wilcoxon signed-rank test, with correction for multiple comparisons within the same metric.

The parameter-mismatch robustness analysis was mainly reported using descriptive curves. For each class of parameter mismatch, changes in performance metrics relative to the no-mismatch condition were calculated at each perturbation level, and were further summarized as the area under the perturbation curve or the maximum performance degradation to quantify controller tolerance to model mismatch. Comparisons of robustness metrics between controllers were likewise performed using within-patient paired tests. Online execution consistency was reported using action consistency, the number of safety-bound violations, and the number of maximum single-step change violations.

All statistical tests were two-sided, with the significance level set at (p<0.05).

S9 Patient recordings and calibration datasets

TABLE S13. Patient recordings and calibration datasets

| **Patient** | **Sex** | **Age at DBS surgery, y** | **Disease duration, y** | **Recording states included** | **Chronic DBS settings at follow-up** |
| --- | --- | --- | --- | --- | --- |
| P1 | M | 59 | 11 | postoperative  MED OFF/DBS ON | L-STN C+6−, 2.40 mA, 70 μs, 140 Hz;  R-STN C+3−, 1.95 mA, 70 μs, 140 Hz |
| P2 | M | 61 | 13 | postoperative  MED OFF/DBS ON | L-STN C+6−, 1.85 mA, 60 μs, 130 Hz;  R-STN C+2−, 2.00 mA, 60 μs, 130 Hz |
| P3 | F | 64 | 9 | postoperative  MED OFF/DBS ON | L-STN C+1−, 2.05 mA, 70 μs, 140 Hz;  R-STN C+5−, 3.10 mA, 70 μs, 140 Hz |
| P4 | F | 56 | 8 | postoperative  MED OFF/DBS ON | L-STN C+3−, 1.45 mA, 70 μs, 130 Hz;  R-STN C+7−, 1.25 mA, 70 μs, 130 Hz |
| P5 | M | 69 | 7 | postoperative  MED OFFDBS ON | L-STN C+6−, 1.40 mA, 70 μs, 130 Hz;  R-STN C+2−, 1.00 mA, 70 μs, 130 Hz |

Patients are anonymized as P1–P5. All patients contributed recordings for patient-specific stimulation–β calibration, stimulation-transition fitting, medication/background modulation estimation, observation-noise estimation, closed-loop simulation, and embedded online validation. P1 additionally provided an independent tuning environment for selecting fixed controller parameters; after parameter selection, all controllers were evaluated with fixed parameters across the five patient-specific models. MED OFF denotes recordings after dopaminergic medication withdrawal, and DBS ON denotes chronic stimulation.
